# Purification-free cryo-EM reveals endogenous assemblies from minute tardigrade samples

**DOI:** 10.64898/2026.10.08.757488

**Authors:** Yohta Fukuda, Daisuke Motooka, Yuko Imamura, Tsuyoshi Inoue

## Abstract

Tardigrades are minute animals renowned for environmental stress tolerance, and their remarkable biology has motivated extensive molecular studies using biochemical and biophysical approaches. Yet the native structures of endogenous macromolecular assemblies in tardigrades remain largely inaccessible, in part because only limited biological material is available. Here we show that purification-free single-particle cryo-EM, guided by long-read cDNA sequence assignment, can recover multiple endogenous assemblies directly from crude extract of a wild-derived *Macrobiotus* cf. *shonaicus* culture. We reconstructed a dimeric vitellogenin (Vg) particle at 3.10 Å resolution, a previously uncharacterized tardigrade ferritin at 3.09 Å resolution, and a low-resolution ribosome map. The Vg particle is a glycosylated and metal-containing assembly with structural evidence of proteolytic processing and contains lipid-like densities occupying a broad chamber whose walls include N-linked glycans. The ferritin structure revealed a divergent C-terminal helix that lines a mineral-like density observed at a position corresponding to previously described mineralization sites in ferritin-family cages. These results establish purification-free cryo-EM as a route to structural analysis of endogenous macromolecular assemblies from minute animal samples. In tardigrades, this approach provides a foundation for future state-dependent structural studies, including desiccation, rehydration and reproduction.

## Introduction

Tardigrades are minute (usually < 1 mm) metazoans renowned for their ability to withstand severe environmental stress, including desiccation^1,2^. This extraordinary tolerance has driven extensive efforts to define tardigrade biology at the molecular level^3^. Much of this work has focused on individual proteins, including Dsup^4^ and abundant heat-soluble protein families^5,6,7,8^, which have been characterized extensively using recombinant proteins, biochemical assays and structural or biophysical approaches. These studies have revealed unusual properties of isolated tardigrade proteins, but they leave largely unexplored the endogenous macromolecular assemblies that exist in animal-derived material. In particular, little is known about the native organization of endogenous macromolecular assemblies in tardigrades, including their maturation states, post-translational modifications, glycans and bound metal ions, because such assemblies are difficult to isolate from minute animals.

Addressing this gap requires methods that can identify and structurally characterize endogenous assemblies without prior purification of a predefined target. This is particularly challenging in minute non-model animals, where the available material is limited and the most informative particles may be heterogeneous, low in abundance or sensitive to biochemical handling. A direct survey of minimally processed material could therefore preserve molecular features that are otherwise lost during purification, while allowing abundant and ordered assemblies to emerge through particle classification. Such an approach would enable structural analysis of endogenous assemblies recovered from animal-derived samples without selecting a target in advance.

Single-particle cryo-EM offers a way to visualize macromolecular assemblies in heterogeneous samples, but most high-resolution applications still rely on biochemical enrichment or purification of a predefined target^9^. Recent studies have begun to relax this requirement by applying lysate-derived material directly to cryo-EM grids. In our previous study using the lysates-are-applied-directly-on-cryo-EM-grids (LApDoG) method, *Escherichia coli* lysate containing a recombinantly expressed tardigrade peroxiredoxin was applied directly to grids, enabling visualization of the protein assembly and its associated nucleic acid-binding states^10^. In parallel, *in extracto* cryo-EM and cryoPRISM analyses of cellular extracts have shown that ribosome-containing complexes can be reconstructed in the presence of cellular components^11,12^. These studies demonstrate the feasibility of atomic-level structural analysis in complex extracts, but have primarily focused on known or readily recognizable targets. We reasoned that, for minute animals, crude extract itself could be used as a structural source and particle classification could reveal which endogenous assemblies are sufficiently abundant and ordered for reconstruction. Such a purification-free approach should be particularly useful when sample amounts are limited, when target identities are not known in advance, or when native states or ligands may be altered or lost during purification.

Here, we applied the LApDoG strategy to crude extract prepared from a wild-derived tardigrade culture. This analysis identified two endogenous storage-related assemblies as major high-resolution reconstructable targets: a dimeric vitellogenin (Vg) particle and a ferritin cage. We also reconstructed a lower-resolution 80S ribosome map and observed two-dimensional classes including proteasome- and catalase-like particles, indicating that the crude extract contained a broader inventory of recognizable endogenous assemblies.

## Results

### Crude-extract cryo-EM captures multiple endogenous assemblies

To test whether endogenous macromolecular assemblies can be structurally recovered directly from minute animal material, we established a laboratory culture from wild tardigrades collected on the campus of The University of Osaka, Suita, Japan (Fig. 1a, Supplementary Fig. 1a). Phylogenetic analysis of the mitochondrial cytochrome *c* oxidase subunit I gene (COI) and nuclear ribosomal internal transcribed spacer 2 (ITS2) placed the culture within the clade containing *Macrobiotus shonaicus*^13^ (Supplementary Fig. 1b-d). Because a detailed morphological diagnosis has not been performed, we refer to the animals as *Macrobiotus cf*. *shonaicus* throughout this study. Crude extract was prepared from approximately 170 animals, including egg-bearing individuals. The animals were starved for 2 days and then homogenized in buffer, followed by centrifugation. The resulting supernatant was applied directly to cryo-EM grids without further biochemical fractionation or target-specific purification (Fig. 1a). Cryo-EM datasets were collected from two grids prepared from the same extract and processed together for the final cryo-EM reconstructions (Supplementary Figs. 2 and 3). Analysis of each grid separately recovered the same major particle types, indicating that the observed assemblies were not restricted to a single grid preparation.

**Fig. 1.**
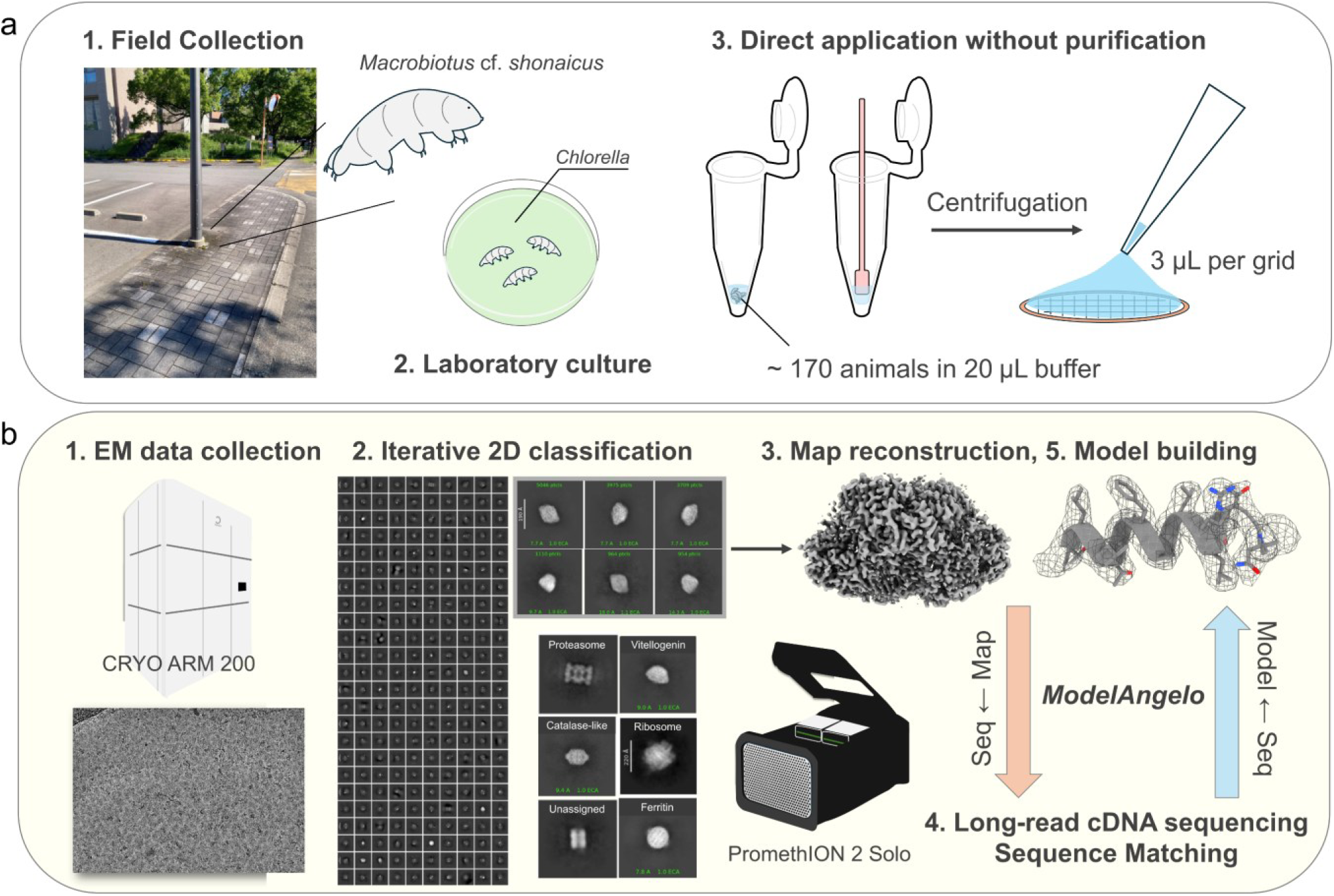
Structure-first cryo-EM analysis of endogenous assemblies from minute tardigrade material without purification. a, Workflow of sample preparation. Wild tardigrades were collected and used to establish a laboratory culture maintained with *Chlorella*. Approximately 170 animals were homogenized in 20 µl of buffer. Following centrifugation, the clarified supernatant was applied directly to each cryo-EM grid without biochemical purification or target-specific enrichment. b, Overview of cryo-EM data collection, particle classification, map reconstruction, long-read cDNA sequence matching and atomic model building. Iterative two-dimensional classification revealed multiple endogenous assemblies. Selected particle classes were subjected to three-dimensional reconstruction. Initial atomic models and partial amino acid sequences were inferred directly from the cryo-EM maps using ModelAngelo. Map-inferred sequences and homologues identified by BLAST were used as references for matching to long-read cDNA sequences obtained from a separate pooled sample from the same laboratory culture. The resulting species-specific amino acid sequences were used for sequence-guided model building and refinement.

The raw micrographs contained a heterogeneous mixture of particles with distinct sizes and morphologies, as expected for unfractionated animal extract (Fig. 1b). Two-dimensional classification nevertheless separated several recognizable classes. Some classes showed features reminiscent of proteasome-like and catalase-like assemblies, although these particles were not pursued to high-resolution reconstruction because of limited particle numbers. Thus, despite the absence of purification, computational classification could resolve multiple macromolecular assemblies from the crude extract.

Three particle classes were sufficient for three-dimensional reconstruction. As described in detail below, the first class yielded a dimeric Vg particle. The second class yielded a previously uncharacterized tardigrade ferritin cage. The third class corresponded to the 80S ribosome (Supplementary Fig. 4a). Three-dimensional reconstruction from 2,257 ribosome particle images yielded a low-resolution (6.57 Å) reconstruction map. Although the ribosome reconstruction did not show sufficient resolution for atomic model building, the 60S subunit was better defined than the 40S subunit (Supplementary Fig. 3f). Rigid-body fitting of human and *Drosophila* reference structures^14,15^ showed that the conserved rRNA and ribosomal protein architecture was broadly accommodated by the map (Supplementary Fig. 4b). We also observed an additional density adjacent to the fitted position of 60S ribosomal protein L29 (Supplementary Fig. 4c). This extra density was not accounted for by either the human or *Drosophila* reference structures. A lineage-specific feature is one possible explanation, but because of the limited resolution of the ribosome map, we did not assign this density.

The recovery of both high-resolution protein assemblies and a lower-resolution ribonucleoprotein complex from the same unfractionated extract demonstrates that crude-extract cryo-EM can capture endogenous particles spanning distinct sizes, compositions and biological functions. The two high-resolution reconstructions provided sufficient map quality for *de novo* model building and sequence inference assisted by ModelAngelo, a machine-learning program for atomic 3D model building in cryo-EM maps^16^. The inferred sequence segments were then compared with transcripts obtained by long-read cDNA sequencing, enabling assignment of the corresponding Vg and ferritin sequences and construction of final atomic models.

### A Vg dimer resolved at 3.10 Å resolution

Among the particle classes recovered from the crude tardigrade extract, one prominent class showed well-defined but initially unassigned 2D projections of an approximately 150-Å particle (Fig. 2a). The class averages displayed oval- to wedge-shaped views, but the identity of the particle was not apparent from the projections alone. *Ab initio* reconstruction with *C*1 symmetry of the selected particles yielded a well-defined three-dimensional map (Supplementary Fig. 2). Subsequent refinement with *C*2 symmetry improved the reconstruction to an overall resolution of 3.10 Å (Fig. 2b). The overall architecture was also recovered in reconstructions without imposed symmetry, supporting the interpretation that the particle represents a genuine dimeric assembly rather than a symmetry-induced reconstruction artifact.

**Fig. 2.**
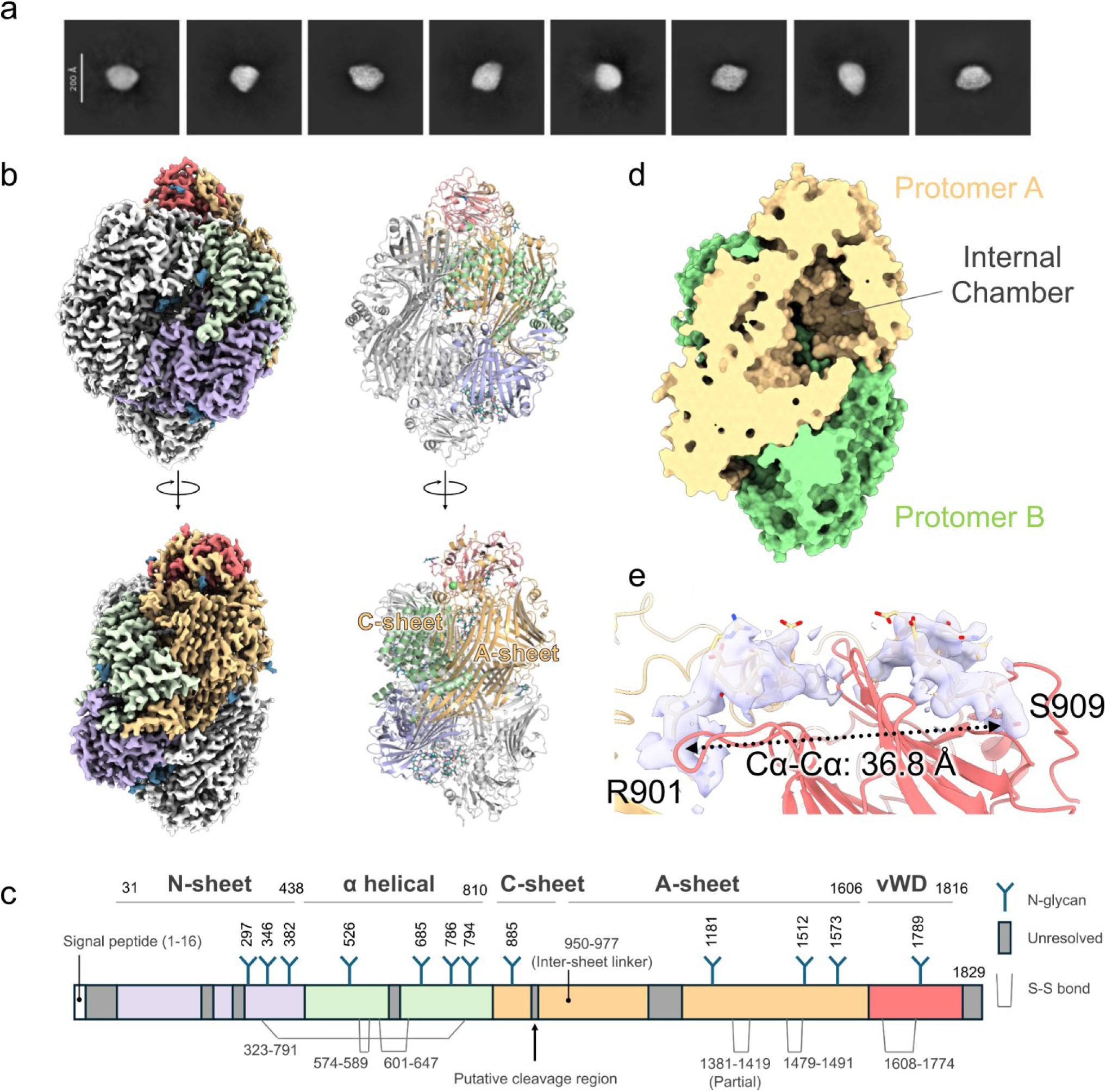
Cryo-EM structure and putative processing of *Ms*Vg6. a, Representative two-dimensional class averages of *Ms*Vg6 particles. Scale bar, 200 Å. b, Cryo-EM map at 3.10 Å resolution (left) and atomic model (right), shown in two orientations. One protomer is colored by domain as in c and the other is gray. c, Organization of the 1,829-residue precursor, showing the N-sheet, α-helical, C-sheet, A-sheet and vWD regions. N-linked glycan sites and disulfide linkages are indicated; unresolved regions are gray. The signal peptide, inter-sheet linker and putative cleavage region are marked. d, Cutaway surface representation showing the internal chamber, with protomers A and B in pale orange and green, respectively. e, Close-up of the putative cleavage region. The Cα atoms of Arg901 and Ser909 are separated by 36.8 Å despite only seven intervening residues in the sequence. The dotted line indicates the measured separation.

The refined map clearly resolved side-chain densities and other local structural features, providing sufficient detail for ModelAngelo-assisted *de novo* model building and sequence inference (Supplementary Fig. 5a, Supplementary File 1). BLAST searches using the inferred sequence segments identified tardigrade Vg6 proteins as the most significant hits, with substantially lower E-values than homologues from non-tardigrade species (Supplementary File 1). Two relatively long segments (245 and 341 residues) gave the same top hit, a vitellogenin 6 (Vg6) like protein from *Paramacrobiotus metropolitanus* (GenBank: XP_055331266.1), with E-values of 7.04 × 10^−67^ and 4.50 × 10^−84^, respectively. Another strong hit was Vg from *Ramazzottius varieornatus* (GenBank: GAU88362.1), whose transcript is highly expressed in active, dehydrated and rehydrated animals^4^. The recovery of a corresponding transcript by long-read cDNA sequencing further supported a tardigrade origin of the particle and enabled construction of the final atomic model. We ultimately assigned the particle as *M*. cf. *shonaicus* Vg6 (*Ms*Vg6).

The final model revealed that the *Ms*Vg6 particle is a homodimer (Fig. 2b). Dimeric assemblies have already been established for Vg-related proteins, including lamprey lipovitellin, whose structure has been determined by X-ray crystallography^17^. The tardigrade particle contains structural features typical for Vg-related proteins: N-sheet, α-helical, C-sheet, A-sheet and von Willebrand factor type D (vWD) regions (Fig. 2b,c). The *Ms*Vg6 protomers enclose a large internal cavity, a characteristic feature of Vg-family proteins (Fig. 2d). The overall fold of an *Ms*Vg6 protomer resembles that of the recently determined cryo-EM structure of full-length honeybee Vg^18^ (Supplementary Fig. 5b).

The reconstructed map of *Ms*Vg6 is unlikely to represent an intact full-length Vg precursor. Although the cDNA-derived sequence encodes an *Ms*Vg6 precursor, the Cα atoms of Arg901 and Ser909 are separated by 36.8 Å despite only seven intervening residues in the sequence (Fig. 2e). Using the standard adjacent Cα–Cα distance of approximately 3.8 Å across a trans peptide bond^50^, the eight intervals between Arg901 and Ser909 give an approximate upper bound of 30.4 Å (8 × 3.8 Å) for the end-to-end distance, shorter than the observed 36.8-Å separation. Moreover, the vWD domain occupies the intervening space, requiring any continuous connection to follow an even longer path around the domain (Fig. 2e). These geometric constraints support a chain discontinuity, provided that the local sequence register is correct. The map is therefore best explained by proteolytic processing of the precursor into a Vg particle composed of discontinuous polypeptide segments.

### N-glycans connect the protomers and line the internal chamber

We next examined non-protein densities associated with *Ms*Vg6. Glycosylation is a common post-translational feature of Vg-family proteins, although individual glycosylation sites are not necessarily conserved across lineages^18,19^. In the cryo-EM map of *Ms*Vg6, N-linked glycan densities were observed at 12 asparagine sites, with the extent of resolved carbohydrate varying among sites (Figs. 2b,c and 3 and Supplementary Fig. 5c). Glycans were modeled where supported by the map. The initial ModelAngelo model assigned nucleic acids to densities at several glycosylation sites that were modeled as N-linked glycans in the final structure (Supplementary Fig. 5a). This may reflect similarities in the local features of glycans and the sugar–phosphate backbone of nucleic acids. The observed N-linked glycans included a high-mannose-type glycan at Asn346, which is located at the dimerization interface (Fig. 3a,b). The N-glycan on Asn346 interacts with another N-glycan on Asn786 from the other protomer (Fig. 3b), suggesting that N-glycans may contribute to stabilization of the dimeric state of *Ms*Vg6. Moreover, the N-linked glycan on Asn786 is positioned along the wall of the internal cavity, where it contributes to the boundary of the chamber (Fig. 3c). Thus, the cavity of *Ms*Vg6 is formed by a composite protein–glycan architecture rather than by protein residues alone. To our knowledge, this is the first structural evidence that N-linked glycans contribute to the wall of a Vg lipid-containing chamber.

**Fig. 3.**
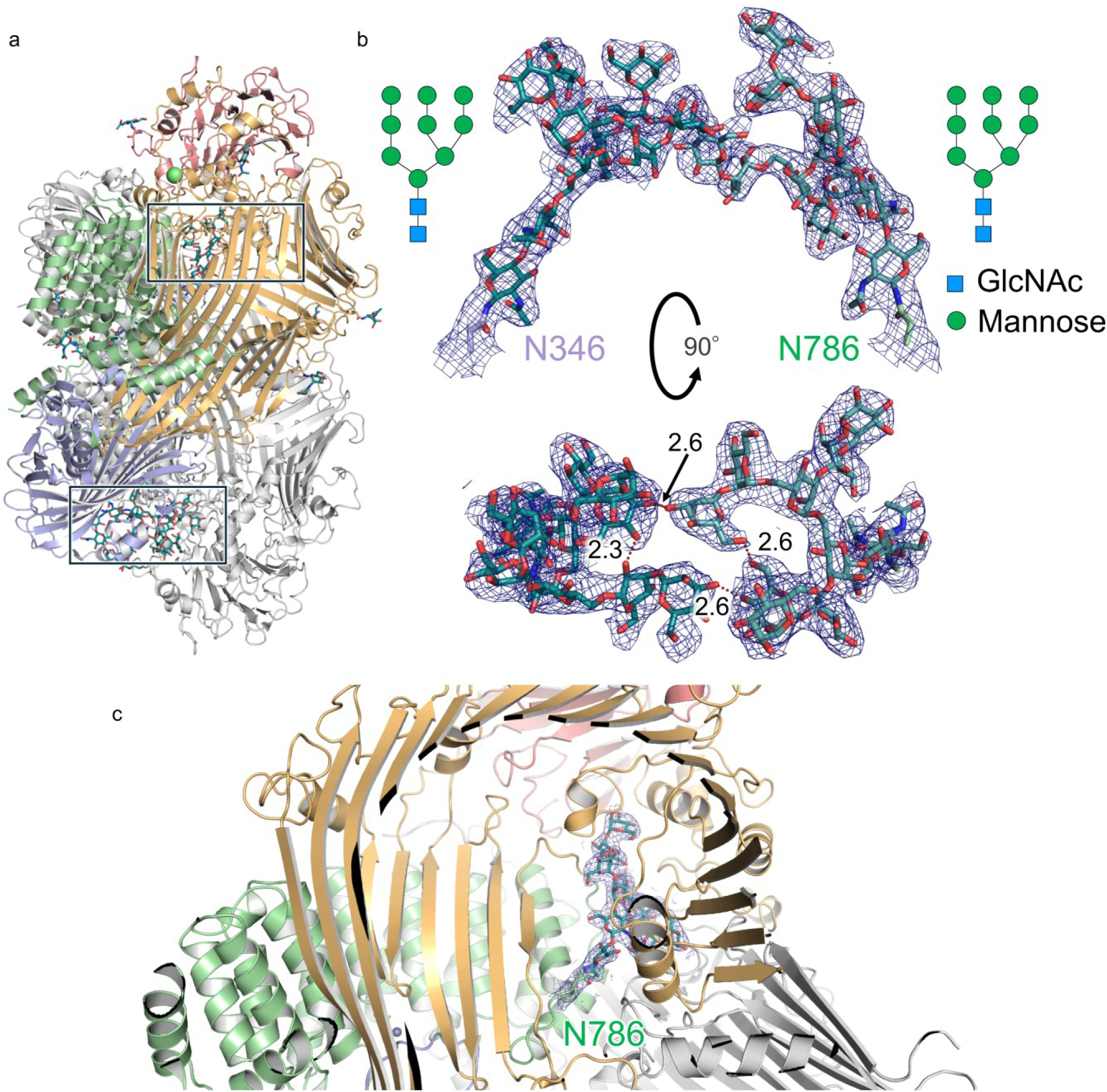
Post-translational modifications by N-glycans in *Ms*Vg6. a, Overview of the dimer with one protomer colored by domain and the other gray. Glycans are shown as sticks. Boxed regions indicate the interprotomer glycan contacts detailed in b. b, Cryo-EM density and models of the N-linked glycans at Asn346 and Asn786 from opposing protomers, shown in two views related by a 90° rotation. Schematics indicate the modeled high-mannose-type glycans. Dotted lines and distances in Å indicate selected interglycan hydrogen-bond contacts. c, Close-up showing the Asn786-linked glycan along the wall of the internal chamber. Cryo-EM density is shown as mesh in b and c.

### Lipid-like densities occupy the internal chamber of *Ms*Vg6

We next examined the densities enclosed within this chamber. Vgs are large yolk precursor lipoproteins that transport and store lipids during reproduction^20,21^. In well-characterized yolk-formation pathways, after uptake into developing oocytes, Vg precursors are proteolytically processed into mature yolk proteins, including lipovitellins, which form lipid-rich storage assemblies^21^. In previously reported crystal and cryo-EM structures of Vg and lipovitellin proteins from other organisms, lipid-associated densities have been observed within large internal cavities or along their inner surfaces^17,18,22,23^. These densities are often heterogeneous or partially ordered rather than being resolved as single well-defined molecules. Consistent with these observations, the cryo-EM map of *Ms*Vg6 revealed multiple elongated non-protein densities within a broad chamber formed by the *Ms*Vg6 protomers and their associated glycans (Figs. 2d and 4a,b,c). Tentative ligand fits illustrate how the elongated densities occupy hydrophobic grooves, whereas polar portions approach charged or polar side chains (Fig. 4b,c). At one site, the fitted ligand lies near residues from both protomers (Fig. 4c, site 1), suggesting that the dimer interface contributes to the cargo-binding environment. These fits do not establish the chemical identities of the ligands or their complete acyl-chain lengths. Nevertheless, the multiple lipid-like densities and their distinct binding environments suggest that *Ms*Vg6 can accommodate diverse lipid cargoes, providing a structural basis for investigating its lipid-storage function.

**Fig. 4.**
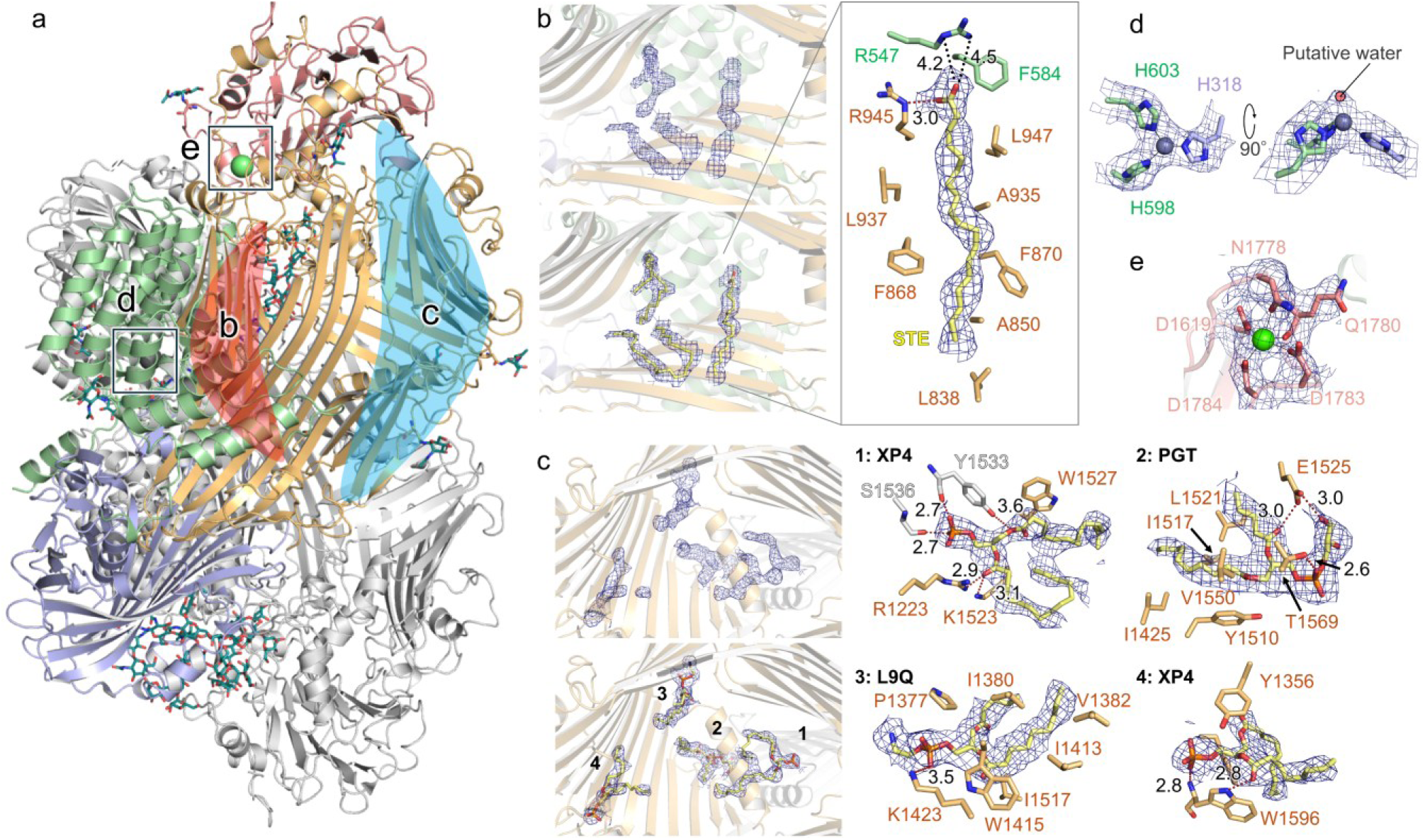
Putative ligand-binding sites in *Ms*Vg6. a, Overview of the dimer, with one protomer colored by domain and the other gray. Putative metal ions are shown as spheres, and glycans and tentative ligand fits are shown as sticks. The indicated regions correspond to b–e. b, Elongated lipid-like density shown without and with a tentative stearic acid fit (PDB chemical component code STE); the enlarged view shows a possible hydrogen bond (3.0 Å) and possible salt bridges (4.2 and 4.5 Å) between the tentatively fitted ligand and surrounding arginine residues. c, Additional lipid-like densities and tentative fits, with four enlarged views numbered 1–4. PDB chemical component codes denote 1,2-dimyristoyl-sn-glycero-3-phosphate (XP4; a phosphatidic acid, sites 1 and 4), phosphatidylglycerol (PGT; site 2) and 1-stearoyl-2-oleoyl-sn-glycero-3-phosphoethanolamine (L9Q; site 3). The indicated distances correspond to possible hydrogen bonds between the tentatively fitted ligands and surrounding residues. d, Putative Zn²⁺ site coordinated by His318, His598 and His603, shown in two views related by a 90° rotation. The adjacent density labeled “Putative water” is consistent with a solvent ligand. e, Putative Ca²⁺ site. Maps are shown as meshes in b–e. Dotted lines and distances in Å indicate selected local contacts. The metal assignments and ligand fits illustrate compatibility with the densities and local environments without establishing elemental identities, definitive ligand chemical identities or complete acyl-chain lengths.

### Putative Zn²⁺- and Ca²⁺-binding environments in *Ms*Vg6

Metal binding has also been reported for Vg-family proteins and lipovitellins, particularly for Zn²⁺ and Ca²⁺, and putative Zn-binding sites in Vg structures often involve histidine-rich environments^18,24,25^. We therefore inspected the cryo-EM map of *Ms*Vg6 for metal-associated densities. The map revealed two types of putative metal-binding site in each protomer. One site is coordinated by His318, His598 and His603 and lies at the interface between the N-sheet and α-helical regions (Fig. 4d). Sequence alignment showed that all three histidines are conserved in the closely related Vgs from *P. metropolitanus* (XP_055331266.1) and *Hypsibius exemplaris* (OQV21904.1) (Supplementary Fig. 6). In the *R*. *varieornatus* homologue (GAU88362.1), the residues corresponding to His318 and His603 are conserved, whereas the position corresponding to His598 is occupied by asparagine. None of these three positions contains histidine in the non-tardigrade homologues included in the alignment, suggesting that the three-histidine metal-binding site may be a lineage-specific feature of a subset of tardigrade Vgs. A small additional density was observed above the putative metal ion at a position compatible with a fourth coordinating ligand, most likely an ordered water molecule or hydroxide ion. Although the elemental identity and protonation state cannot be determined from the cryo-EM map alone, this 3His-solvent coordination environment is characteristic of many Zn^2+^-binding sites, including the canonical zinc center of carbonic anhydrase, in which Zn^2+^ is tetrahedrally coordinated by three histidine residues and a water/hydroxide ligand^26^. We therefore modeled a Zn^2+^ ion at this putative metal-binding site.

We also observed an elongated density near the putative Zn site that could accommodate a sphingosine-like molecule (Supplementary Fig. 7a). Sphingosine is a product of ceramide hydrolysis by ceramidases.^27^ Comparison of the local metal-binding environments of *Ms*Vg6 and human alkaline ceramidase 3 (ACER3; PDB 6G7O)^27^ highlighted a shared three-histidine coordination arrangement and nearby serine and acidic residues (Supplementary Fig. 7a,b). In *Ms*Vg6, Glu718, Ser321 and Ser714 surround the region above the putative Zn site. In ACER3, Ser77 and Asp92 form hydrogen bonds with the Zn-bound water molecule,^27^ and mutational analysis has established their importance for ceramide hydrolysis.^28^ Sequence comparison further showed that, in the *P. metropolitanus* homologue (XP_055331266.1), the positions corresponding to Ser321 and Glu718 are occupied by threonine and aspartate, respectively, preserving a hydroxyl-bearing side chain and an acidic side chain at these positions. In the *H. exemplaris* homologue (OQV21904.1), Ser321 is conserved, whereas Glu718 is replaced by lysine, indicating a difference in the local chemical environment. Ser714 is not conserved in either homologue (Supplementary Fig. 6). The sphingosine-like density and these local similarities raise the possibility that the putative Zn site contributes to lipid hydrolysis within the chamber. However, the density could also represent a bound lipid acquired independently of catalysis. The proposed catalytic role therefore remains a testable hypothesis, as neither the ligand identity nor ceramidase activity of *Ms*Vg6 has been established.

The other putative metal site was observed at a distinct position and showed a coordination environment compatible with a Ca²⁺ ion (Fig. 4e). The site is surrounded by Asp1619, Asn1778, Gln1780, Asp1783 and Asp1784 in the vWD region and is spatially separated from the putative Zn site. Potential oxygen-donor residues are retained at these positions in the six aligned tardigrade homologues from three other species (Supplementary Fig. 6). Furthermore, structural superposition with honeybee Vg (PDB 9ENR) revealed a similar local arrangement of oxygen-containing side chains at the corresponding site (Supplementary Fig. 7c), consistent with conservation of the putative metal-binding environment despite differences in individual residue identities (Supplementary Fig. 6). In honeybee Vg, Ca²⁺ binding at this site was predicted using AlphaFold 3, but the ion was not modeled in the experimental cryo-EM structure.^18^ The density observed at the corresponding site in *Ms*Vg6 therefore provides experimental structural support for metal binding at a vWD site shared by tardigrade and honeybee Vgs. Because no independent elemental analysis was performed, we refer to this site as a putative Ca²⁺-binding site.

Collectively the lipid-like densities, N-linked glycans and putative metal-binding sites show that the *Ms*Vg6 particle retains multiple non-polypeptide features in the crude tardigrade extract, indicating that purification-free cryo-EM recovered not merely a Vg-family protein fold, but an endogenous assembly with resolved post-translational structural features. These features provide a structural basis for elucidating the functions of *Ms*Vg6.

### A new tardigrade ferritin with an extended C terminus

Another abundant particle class recovered from the crude tardigrade extract showed spherical projections (Fig. 5a). Ab initio reconstruction and three-dimensional refinement yielded a hollow, nearly spherical cage with octahedral symmetry, which was refined to 3.09 Å resolution (Fig. 5b). The map showed clear secondary-structure and side-chain densities, allowing *de novo* model building and sequence inference (Supplementary Fig. 8). BLAST searches using a 189-residue consensus sequence derived from ModelAngelo predictions identified ferritin-family proteins, with a soma ferritin-like protein from *Paramacrobiotus metropolitanus* (GenBank: XP_055355121.1) as the top hit (41.6% sequence identity, 78.8% query coverage; E-value, 7.61 × 10^−29^; Supplementary File 2). Comparison of the consensus sequence with long-read cDNA-derived transcripts identified a corresponding ferritin-like transcript from the *M*. cf. *shonaicus* culture (*Ms*Ftn), supporting assignment of the particle as an endogenous tardigrade ferritin. SignalP 6.0^51^ predicted an N-terminal signal peptide in *Ms*Ftn (Supplementary Table 1), consistent with entry into the secretory pathway.

**Fig. 5.**
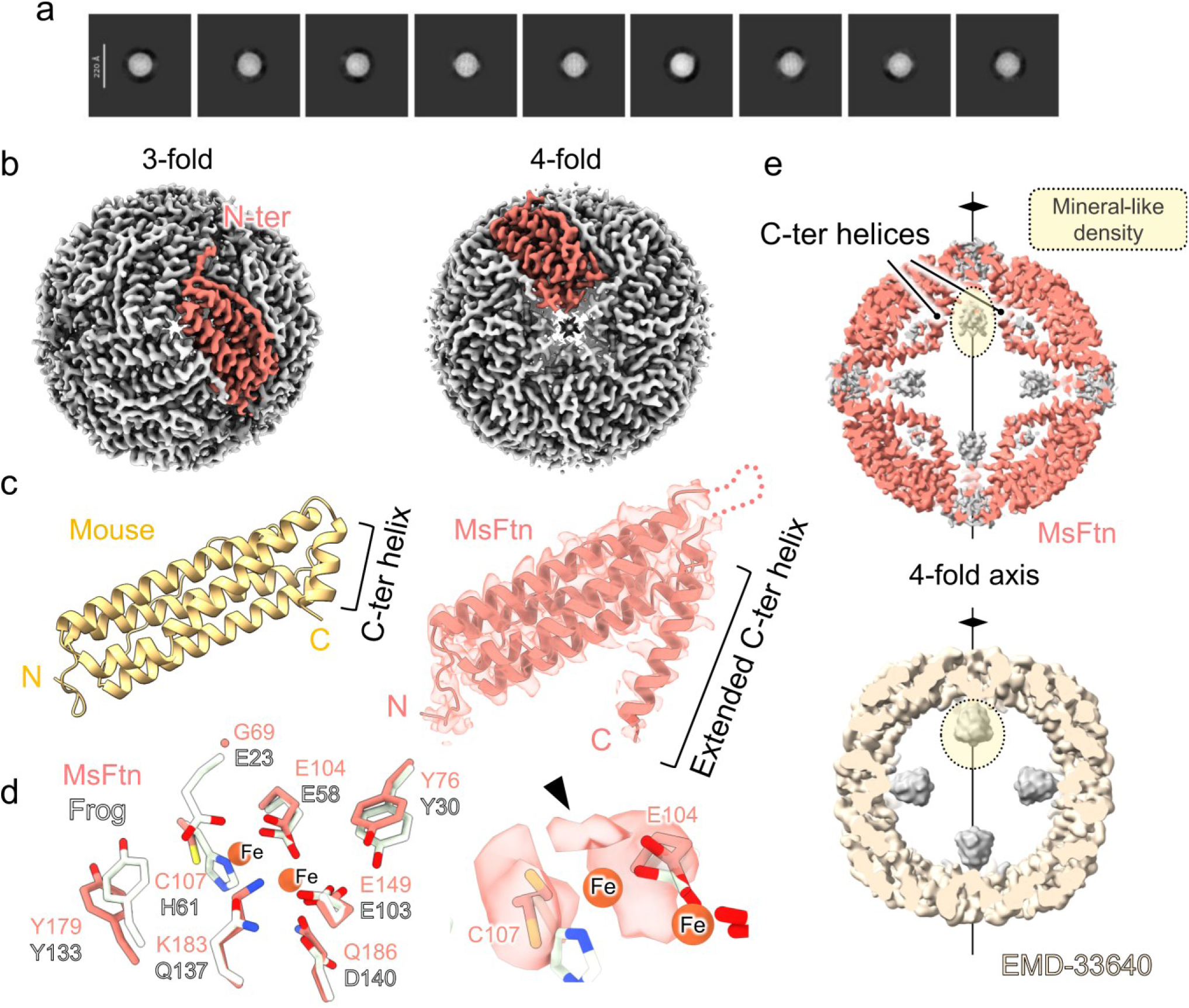
*Ms*Ftn cage architecture and mineral-associated density. a, Representative two-dimensional class averages of *Ms*Ftn particles. Scale bar, 220 Å. b, Cryo-EM reconstruction at 3.09 Å resolution viewed along the threefold and fourfold axes, with a representative subunit highlighted in salmon. c, Comparison of a mouse heavy-chain ferritin protomer (yellow; PDB 7A4M)^31^ and *Ms*Ftn (salmon), highlighting the extended C-terminal helix of *Ms*Ftn. d, Comparison of the putative ferroxidase-center region of *Ms*Ftn (salmon) with bullfrog M ferritin (white; PDB 3RBC)^30^. Residues are labeled in the corresponding colors. Orange spheres show the iron positions in the reference structure. The enlarged view shows *Ms*Ftn map density near the corresponding site (arrowhead), together with Glu104 and Cys107. The cryo-EM map indicates that Cys107 adopts alternative side-chain conformations. e, Sectioned maps showing the luminal region near the fourfold axis of *Ms*Ftn (upper) and *Streptomyces coelicolor* bacterioferritin (lower; EMD-33640)^32^. Dotted outlines highlight mineral-like density, and the *Ms*Ftn C-terminal helices are indicated.

Ferritins are protein cages conserved from bacteria to eukaryotes^29^. They regulate intracellular iron by oxidizing Fe^2+^ at ferroxidase centers and sequestering the resulting Fe^3+^ as a ferric mineral within an internal cavity, thereby contributing to iron storage and detoxification. This chemistry enables ferritins to store iron, limit iron-mediated oxidative damage and provide an iron reserve for cellular metabolism. *Ms*Ftn assembles into a 24-subunit cage composed of protomers with the canonical ferritin four-helix-bundle fold (Fig. 5b,c). The subunits pack around the threefold and fourfold symmetry axes to generate an internal lumen, and the overall cage architecture is closely similar to those of other ferritin-family proteins despite limited sequence conservation. Within each protomer, the region corresponding structurally to the ferroxidase center lies within the helical bundle, although the canonical metal-coordinating residues are not fully conserved (Fig. 5d, Supplementary Fig. 9). In the comparison with bullfrog M ferritin containing two iron ions at the ferroxidase center (PDB 3RBC)^30^, Glu104 and Glu149 of *Ms*Ftn correspond to the iron-coordinating residues Glu58 and Glu103 in bullfrog M ferritin, respectively. Tyr76 and Tyr179 also correspond to Tyr30 and Tyr133 in bullfrog M ferritin. By contrast, Glu23, His61, Gln137 and Asp140 in bullfrog M ferritin correspond to Gly69, Cys107, Lys183 and Gln186 in *Ms*Ftn, respectively. In particular, the side chain of Lys183 may interfere with metal binding in the canonical configuration. Nevertheless, a weak spherical density was observed at one position within this putative ferroxidase center, compatible with a metal ion at this site (Fig. 5d). Although the elemental identity cannot be determined from the cryo-EM map alone, the density and surrounding residues suggest a potential metal-binding site involving Glu104, Cys107 and Glu149. Ferroxidase activity remains to be established.

Comparison with canonical eukaryotic ferritin structures revealed a distinctive feature of *Ms*Ftn: an extended C-terminal helix on the luminal side of the cage (Fig. 5c, Supplementary Fig. 9). For example, in mouse heavy-chain ferritin (PDB 7A4M)^31^, the corresponding C-terminal region is shorter and does not form an equivalent luminal extension. Thus, *Ms*Ftn preserves the conserved ferritin cage architecture while modifying the luminal surface through the extended C-terminal helix.

These observations identify the second high-resolution assembly as a tardigrade ferritin distinct from known ferritin family proteins. The structure combines a canonical ferritin cage with an ordered luminal C-terminal helix, providing a structural framework for examining how *Ms*Ftn may function. Further biochemical implications obtained from our *Ms*Ftn structure are discussed below.

### Ferritin internal density suggests a mineral-associated state

We next examined the internal cavity of *Ms*Ftn, focusing on non-protein densities associated with the luminal surface and symmetry-axis regions of the cage. The cryo-EM map contained a prominent internal density on the luminal side of the fourfold channel (Fig. 5e). This density was spatially separated from the modeled protein backbone and could not be assigned to *Ms*Ftn. It was therefore left unmodeled and provisionally interpreted as mineral-like density within the ferritin lumen. A corresponding internal density was also observed in the reconstruction obtained without imposed symmetry (*C*1), supporting that this feature is not an artifact of octahedral symmetry imposition (Supplementary Fig. 10a).

The position of this density was notable. Rather than forming a diffuse core at the center of the cage, the density was localized beneath the fourfold-axis region, where ferritin-family cages provide a highly symmetric internal environment^29^. A similar fourfold-axis mineral-associated density has been reported in *Streptomyces coelicolor* bacterioferritin, where it was interpreted as an early iron-mineralization state^32^ (Fig. 5e). Comparison of cryo-EM maps showed that the tardigrade ferritin density occupies a corresponding luminal position beneath the fourfold channel. This positional correspondence supports the interpretation that the cryo-EM density in the *Ms*Ftn cavity is mineral-associated, although its chemical composition cannot be determined from the cryo-EM map alone.

The extended C-terminal region of *Ms*Ftn is positioned adjacent to this internal density (Fig. 5e). The segment follows the luminal surface near the fourfold-axis region and runs along the mineral-like density. Thus, the C-terminal extension is not simply an appended terminal feature, but forms part of the local luminal architecture surrounding the putative mineralization site. However, the functional role of this extension cannot be inferred from the structure alone. Future biochemical and mutational analyses will be needed to determine whether this C-terminal helix contributes to iron uptake, mineral nucleation, mineral retention or release from the ferritin cage.

We also observed an additional elongated density along the inner wall of the cage (Supplementary Fig. 10b). Although the C terminus is oriented toward the lumen, the cDNA-derived sequence model indicates that the ordered C-terminal segment is not long enough to account for this separate density without disrupting the fitted protein model. We therefore left this feature unmodeled and refer to it as an unidentified inner-wall density. Its presence suggests that the endogenous ferritin cage may contain additional lumen-associated components or mineral-related density that are not captured by the protein sequence alone. The fourfold-axis mineral-like density suggests that the tardigrade ferritin was captured in an endogenous mineral-associated state, providing a structural snapshot consistent with its iron-storage function in the animal. Importantly, this state was recovered directly from crude animal extract without *in vitro* iron loading or reconstitution.

## Discussion

Understanding how proteins function in organisms requires structural information about their endogenous assemblies and associated molecular components. Here, crude extract from approximately 170 tardigrades yielded high-resolution structures of two previously uncharacterized endogenous assemblies without target-specific purification. Combining map-based sequence inference with long-read cDNA sequencing linked these structures to their encoding transcripts without a pre-existing genome assembly. This makes the approach useful for discovering the molecular organization of non-model animals: assemblies can first be recognized by their structures, assigned to sequences and then examined for features that suggest biological functions. These reconstructions were obtained using a 200-kV cryo-electron microscope, demonstrating that structural analysis of these minute animal samples was feasible at this accelerating voltage. The recovery of an 80S ribosome and additional recognizable particle classes from the same extract further shows that a single preparation can provide access to multiple endogenous complexes.

The structures reveal features beyond the protein scaffolds. In *Ms*Vg6, N-linked glycans participate in contacts between protomers and form part of the boundary of a chamber containing lipid-like densities. Dimerization and the presence of N-glycans and lipids in the inner chamber themselves have precedents in lipovitellin structures^17,18^; however, the present structure reveals how glycans, the processed polypeptide architecture and associated non-protein densities are organized together in a tardigrade assembly. The glycan arrangement suggests that glycosylation could influence both dimer stability and the cargo-binding environment, providing specific sites for functional analysis. The putative Zn-binding site raises a further question: could this chamber support lipid chemistry as well as storage? The three coordinating histidines are conserved in some tardigrade Vgs. This coordination environment, nearby polar residues and adjacent sphingosine-like density motivate this possibility, although ligand identity and catalytic activity remain unresolved. By contrast, the putative Ca-binding environment is shared with honeybee Vg^18^. These observations distinguish potentially lineage-specific features from conserved ones and provide a structural basis for investigating their respective roles.

*Ms*Ftn illustrates how a conserved cage architecture can accommodate a different internal environment. Its extended C-terminal helix lines the lumen adjacent to mineral-like density, while substitutions at positions corresponding to the canonical ferroxidase center alter the potential metal-binding chemistry. The coexistence of these features makes it important to test how iron oxidation and mineral accumulation are coupled in this ferritin, rather than assuming that a conserved overall fold implies an unchanged mechanism. The mineral-like density occupies a position comparable to that reported in bacterioferritin^32^ and remains visible without imposed symmetry. Although its composition is not established, its recovery without in vitro iron loading provides an endogenous structural context for testing whether the C-terminal extension contributes to mineral nucleation or retention. This context would be difficult to infer from the protein sequence or an unliganded cage alone.

The scope of this approach is defined by what survives extraction and can be reconstructed. Particle abundance, size, stability and conformational order influence which assemblies are recovered^33,34^, and the present preparation principally samples soluble or extractable material. It therefore does not provide a complete inventory of the animal’s macromolecular complexes. Nor does extraction preserve cellular localization or establish that every observed interaction existed unchanged in the animal. In particular, the timing of *Ms*Vg6 processing remains unresolved. Within these limits, the retained glycans, lipid-like densities and mineral-like density demonstrate access to molecular features relevant to function, while directing biochemical work toward specific sites and interactions.

A next application is to ask how these endogenous assemblies change with physiological state. In tardigrades, comparisons across reproduction, fasting, desiccation and rehydration could test whether glycan-mediated contacts, cargo occupancy or ferritin mineral-associated densities change alongside organismal responses. The present structures do not establish a role in stress tolerance, but they identify concrete molecular features through which such roles can be investigated. More broadly, coupling structural discovery to sequence assignment allows work on poorly characterized organisms to begin with the assemblies and associated components recovered from the organism itself, and to use that information to formulate targeted functional experiments.

## Methods

### Tardigrade collection, culture and species assignment

Tardigrades were collected from dried moss on the Suita Campus of The University of Osaka, Osaka, Japan. A laboratory culture was established from the wild-collected animals. The culture was maintained on agarose plates supplemented with Volvic water and fed with *Chlorella* (CHIKUGO [CK-5]; Chlorella Industry Co., Ltd.). Animals used for cryo-EM sample preparation were taken from this laboratory culture. Species assignment was based on molecular barcoding of mitochondrial cytochrome *c* oxidase subunit I (COI) and nuclear ribosomal internal transcribed spacer 2 (ITS2). Before DNA preparation, a single animal was collected from a culture plate and starved for 2 days on agarose plates without *Chlorella*. After starvation, the animal was washed twice with Volvic water and then twice with ultrapure water to reduce carryover of food algae and external contaminants. Genomic DNA was prepared by alkaline lysis using a HotSHOT method^35^ adapted for tardigrades^36^. Briefly, the washed animal was transferred to a PCR tube containing alkaline lysis buffer (25 mM NaOH, 0.2 mM EDTA) and incubated at 95 °C for 10 min. The lysate was then neutralized by addition of neutralization buffer (40 mM Tris-HCl, pH 8.0) and used directly as the template for PCR amplification of both COI and ITS2 fragments from the same individual. PCR was performed for 40 cycles of denaturation at 98 °C for 10 s, annealing at 42 °C for 5 s and extension at 68 °C for 15 s, using KOD One PCR Master Mix (TOYOBO) and previously reported primer sets (refs. 37,38) listed in Supplementary Table 1. Amplified fragments were analyzed by agarose gel electrophoresis, and bands of the expected sizes were excised and purified using the NucleoSpin Gel and PCR Clean-up kit (MACHEREY-NAGEL). Purified PCR products were subjected to Sanger sequencing by FASMAC Co., Ltd. Partial COI and ITS2 sequences corresponding to high-confidence regions of the Sanger reads obtained from the PCR amplicons were used for subsequent analyses and shown in Supplementary Table 1. These partial sequences were initially queried against the NCBI nucleotide database using BLASTn.^39^ Because the closest matches were sequences from tardigrades, subsequent phylogenetic analyses were performed within Tardigrada. For the broad COI analysis, representatives of major lineages within Macrobiotidae and selected other eutardigrade groups were retrieved from GenBank using the rentrez package.^40^ Reference sequences of *Macrobiotus shonaicus* and related *Macrobiotus* species were additionally collected for focused COI and ITS2 analyses of close relatives. Sequences were aligned using DECIPHER^41^. After alignment, each dataset was trimmed to query-anchored columns corresponding to non-gap positions in the sequence of the present sample. Regions of 510 bp, 621 bp and 307 bp were retained for the broad COI, focused COI and focused ITS2 analyses, respectively. Maximum-likelihood trees were inferred in phangorn^42^ under a GTR+I+G4 model with 1000 bootstrap replicates. The broad COI tree was rooted with *Milnesium tardigradum*, whereas the close-relative COI and ITS2 trees were rooted with *Mesobiotus harmsworthi*. Trees were visualized using the Interactive Tree Of Life (iTOL) web tool.^43^

### Long-read cDNA sequencing and sequence assignment

Total RNA was extracted from a pooled sample of 265 animals from the same *M*. cf. *shonaicus* laboratory culture used for cryo-EM analysis. Animals were collected from culture plates and starved for 2 days on agarose plates without *Chlorella*. After starvation, the animals were transferred to a clean 1.5-ml tube and washed twice with Volvic water and then twice with ultrapure water to reduce carryover of food algae and external contaminants. After the final wash, residual water surrounding the animals was removed as completely as possible by pipetting carefully while retaining the animals in the tube. Total RNA was then extracted using the Direct-zol RNA MiniPrep kit (Zymo Research) according to the manufacturer’s protocol, including on-column DNase treatment.

Full-length cDNA was synthesized from total RNA using the SMARTer PCR cDNA Synthesis Kit (Takara Bio, Shiga, Japan). Sequencing libraries were prepared using the Ligation Sequencing Kit V14 (SQK-LSK114; Oxford Nanopore Technologies) and sequenced on a PromethION P2 Solo system with a FLO-PRO114M flow cell. Base calling was performed with Dorado v1.0.0 using the super-accuracy (SUP) model. Candidate cDNA reads corresponding to the Vg6 and ferritin-like proteins observed in the cryo-EM maps were identified using NCBI BLAST+ v2.17.0.^39^ For protein-family identification, map-inferred Vg sequence segments (245 and 341 residues), and a 189-residue ferritin consensus sequence, were queried against the NCBI clustered_nr database using BLASTp (Supplementary Files 1 and 2). For the Vg6 protein, the amino acid sequence initially inferred by ModelAngelo was used to identify a homologous Vg6-like protein from *H*. *exemplaris* by BLAST searching. Although the *P. metropolitanus* sequence was the top database hit, it was annotated as low quality. The *H. exemplaris* Vg6-like sequence (GenBank: OQV21904.1) was therefore used as the protein reference for local blastx searches of the long-read cDNA dataset. For the ferritin-like protein, the ModelAngelo-derived amino acid sequence was used directly as the protein reference for blastx searches.

Initial Vg6-like candidate reads were identified by blastx against the *H*. *exemplaris* Vg6-like protein and were then recovered by nucleotide-level searches using cDNA-derived baits. The first 2,000,000 reads were screened. Near-full-length Vg6-like candidates were required to be 5,000–6,500 nt long, to align over at least 5,000 nt and to cover at least 85% of the best nucleotide bait. Ferritin-like candidates were identified from the first 200,000 reads by blastx against the 189-residue map-derived ferritin sequence. Reads used for the initial ferritin consensus were required to cover at least 95% of the target in a single orientation, with one HSP of at least 180 amino acids, at least 65% amino acid identity and a read length of 700–1,600 nt.

Candidate reads were oriented according to their BLAST alignments and compared by all-versus-all blastn. For these comparisons, pairwise nucleotide identity was taken from the highest-bit-score HSP for each read pair. Coverage was defined as the aligned span divided by the corresponding bait or target length. Closely related near-full-length reads were grouped by average-linkage hierarchical clustering using a distance of 1 − identity/100. Consensus construction was restricted to a conservatively selected subset of near-full-length reads with consistent orientation, extensive target coverage and high pairwise nucleotide identity within each selected cluster. The medoid was defined as the read with the highest mean pairwise nucleotide identity to the other reads in the selected cluster. The selected reads were aligned using AlignSeqs in DECIPHER.^41^ In each multiple-sequence alignment, a nucleotide or gap was called only when supported by a strict majority of the selected reads. Majority-supported gap columns were removed. At positions lacking a strict majority, the nucleotide from the medoid read was used and flagged in the analysis output. These medoid-derived calls were retained in the final cDNA sequences, which were not altered on the basis of the cryo-EM maps. No independent nucleotide-level validation was performed. Open reading frames were identified in the resulting consensus cDNA sequences and translated using Biostrings. The resulting protein sequences were used for sequence-guided model building. Read identifiers are provided in Supplementary Table 3. The final coding and amino acid sequences are provided in Supplementary Table 1.

N-terminal signal peptides were predicted from the cDNA-derived amino acid sequences of *Ms*Vg6 and *Ms*Ftn using SignalP 6.0^51^ (Supplementary Table 1). The *Ms*Ftn sequence contains two candidate translation-initiation ATG codons, with the downstream candidate corresponding to Met18 under the numbering used here. The upstream candidate was used to define Met1 in the reported coding and amino acid sequences and for residue numbering throughout this study. The translation-initiation site was not experimentally determined.

### Preparation of crude tardigrade extract

For cryo-EM sample preparation, approximately 170 animals were starved for 2 days before extraction. The animals were washed twice with Volvic water and then twice with 10 mM phosphate buffer, pH 7.0. The washed animals were transferred to 20 µL of 10 mM phosphate buffer, pH 7.0, and disrupted manually with a pestle on ice. The homogenate was centrifuged at 10,000 × g for 15 min at 4 °C, and the resulting supernatant was used directly for cryo-EM grid preparation without further biochemical fractionation, affinity capture or target-specific purification. The supernatant had an orange color after extraction. No protease inhibitors were used, and the extract was vitrified within 1 h of extraction.

### Cryo-EM grid preparation and data collection

A 3-µL aliquot of the crude tardigrade extract was applied to QUANTIFOIL R1.2/1.3 Cu 200 mesh grids (Quantifoil Micro Tools GmbH) that had been freshly glow-discharged at 20 mA for 20 s using a JEC-3000FC Auto Fine Coater (JEOL). Grids were blotted and plunge-frozen in liquid ethane using a Vitrobot Mark IV (Thermo Fisher Scientific) operated at 8 °C and 100% humidity, with a blot time of 3 s, a blot force of −10 and a wait time of 5 s. Cryo-EM movies were collected from two grids prepared from the same crude extract. Grid 1 yielded 10,591 movies and grid 2 yielded 8,416 movies. Data were collected on a CRYO ARM 200 (JEOL) operated at 200 kV and equipped with a K3 direct electron detector (Gatan). Movies were recorded using SerialEM^44^ at a nominal magnification of 60,000, corresponding to a calibrated pixel size of 0.83 Å per pixel, with a total electron exposure of 40 e⁻ Å⁻^2^ fractionated over 40 frames. The nominal defocus range was −0.7 to −1.7 µm. The image-processing workflow is shown in Supplementary Figs. 2 and 3, and data collection, reconstruction and refinement statistics are summarized in Supplementary Table 2.

### Cryo-EM image processing

Cryo-EM image processing was performed using CryoSPARC ver 4.7.1.^45^ Movie frames were aligned and dose-weighted by patch motion correction, and contrast transfer function parameters were estimated by patch CTF estimation. Exposures with poor CTF fits, excessive motion, crystalline ice or severe contamination were removed during exposure curation.

Initial particles were picked from a subset of micrographs using blob picking and subjected to iterative two-dimensional classification. Representative class averages corresponding to recognizable particle types were used as templates for template-based particle picking from the full dataset. The resulting particles were subjected to iterative two-dimensional classification to separate distinct endogenous assemblies. Classes corresponding to *Ms*Vg6, *Ms*Ftn and the 80S ribosome were processed independently. Additional classes with proteasome-like and catalase-like features were identified but were not pursued to high-resolution reconstruction because of limited particle numbers.

For the *Ms*Vg6 particle, selected particles were used for ab initio reconstruction without imposed symmetry. The resulting map revealed a dimeric architecture. Subsequent heterogeneous refinement, non-uniform refinement (using *C*2 symmetry), global/local CTF refinement, and reference-based motion correction were performed. Reconstructions without imposed symmetry were also examined to confirm that the dimeric architecture was not an artifact of imposed symmetry. Further three-dimensional classification to improve the homogeneity of lipid-like density selected 17,127 particles, which yielded the final 3.10-Å reconstruction after non-uniform refinement with *C*2 symmetry, based on the gold-standard Fourier shell correlation criterion (Supplementary Fig. 2a). This final map was used for all *Ms*Vg6 map visualizations in the figures.

For *Ms*Ftn, particles were selected after iterative two-dimensional classification. Ab initio reconstruction and three-dimensional refinement yielded a ferritin-like cage. Subsequent non-uniform refinement with O symmetry and global CTF refinement were performed, and map handedness was corrected (Supplementary Fig. 3a). Final non-uniform refinement was performed with octahedral symmetry imposed, resulting in a 3.09-Å reconstruction. Reconstructions with *C*1, *C*3 and *C*4 symmetry were also inspected (Supplementary Fig. 3a). The *C*1 reconstruction was used to assess whether the mineral-like internal density was retained without imposed symmetry (Supplementary Fig. 10a).

A third particle class corresponding to the 80S ribosome was reconstructed through ab initio reconstruction, heterogeneous and homogeneous refinement, and final non-uniform refinement with *C*1 symmetry, yielding a 6.57-Å map from 2,257 particles (Supplementary Fig. 3e). This map was used to assess the ability of the crude-extract dataset to recover a large endogenous ribonucleoprotein complex, but was not used for atomic model building. The human (PDB 4UG0) and *Drosophila* (PDB 4V6W) reference models were independently fitted as whole rigid bodies into the map using UCSF ChimeraX. After fitting, map regions within 8 Å of each reference model were colored according to that model for comparison (Supplementary Fig. 4).

### Atomic model building and structure refinement

Initial atomic models of *Ms*Vg6 and *Ms*Ftn were generated from the cryo-EM maps using ModelAngelo^16^. Amino acid sequence segments inferred from the maps were compared with cDNA transcripts obtained from the same laboratory culture as described in the “Long-read cDNA sequencing and sequence assignment” section. The primary consensus protein sequences obtained from the cDNA analysis were supplied to ModelAngelo through RELION v5.0 for sequence-guided model building against the corresponding cryo-EM maps. The ModelAngelo-generated models were inspected and manually corrected in Coot^46^.

For *Ms*Vg6, N-linked glycan residues were added at asparagine sites for which the cryo-EM map showed unambiguous branched carbohydrate density. Glycan composition and connectivity were modeled conservatively according to the resolved density, and unresolved distal carbohydrate moieties were not included. Lipid-like densities within the internal cavity of *Ms*Vg6 were not assigned definitive chemical identities. Tentative ligand fits are shown in Fig. 4b,c and Supplementary Fig. 7a to illustrate their compatibility with the density and local contacts; these fits do not establish the lipid species or complete acyl-chain lengths. The tentative ligand models are included in the deposited coordinates to facilitate reproduction of the figures, without implying definitive chemical identification. The two types of putative metal-binding site in each *Ms*Vg6 protomer were modeled on the basis of their coordination environments. At the histidine-rich site, a Zn²⁺ ion was modeled on the basis of the density and three-histidine coordination geometry. Adjacent density was interpreted as a possible solvent ligand but was not included as a solvent molecule in the atomic model. A second site was modeled as a putative Ca²⁺ ion based on its local coordination geometry. Because elemental identities could not be established directly from the cryo-EM map, these assignments were treated as tentative. For *Ms*Ftn, the prominent fourfold-axis mineral-like density and the additional inner-wall density were not assigned to specific chemical species and were left unmodeled. Real-space refinement of the models against the corresponding cryo-EM maps was performed using PHENIX^47^. Model geometry was evaluated using MolProbity.^48^

Protein sequences were aligned using Clustal Omega, with residue numbering referenced to *Ms*Vg6 or *Ms*Ftn. The alignments were visualized using ESPript^49^ (Supplementary Figs. 6 and 9).

## Data availability

The cryo-EM density maps and the atomic coordinates of *Ms*Vg6 and *Ms*Ftn have been deposited in the Electron Microscopy Data Bank (EMDB) and the PDB with EMDB accession codes EMD-XXXX and EMD-YYYY, and PDB accession codes XXXX and YYYY, respectively. The cryo-EM density map of the 80S ribosome has been deposited in the EMDB with the EMDB accession code EMD-ZZZZ. The Nanopore cDNA sequencing reads are deposited in the DDBJ Sequence Read Archive (DRA) under experiment accession DRX1175545 and run accession DRR1206612, linked to BioProject PRJDB45826 and BioSample SAMD02113280. The final coding and amino acid sequences of *Ms*Vg6 and *Ms*Ftn and the partial COI and ITS2 sequences used for species assignment are provided in Supplementary Table 1.

## Acknowledgements

We thank Mr. Hiroki Tanino and Ms. Yuki Shimokawa (Graduate School of Pharmaceutical Sciences, the University of Osaka) for their support in cryo-EM data collection. The authors used ChatGPT to assist with English language editing and grammatical improvement. All AI-assisted text was reviewed and edited by the authors, who take full responsibility for the final content of the manuscript.

## Author contributions

Y.F. conceptualized the project, collected tardigrades, established the tardigrade culture, prepared total RNA from tardigrades, performed bioinformatics analyses, prepared the cryo-EM samples, collected the cryo-EM data, processed the cryo-EM data, modeled the protein structures and wrote the manuscript. D.M. and Y.I. prepared cDNA from tardigrade RNA and performed cDNA sequencing. T.I. supervised the study. All authors approved the final version of the manuscript.

---

**Supplementary Fig. 1.**
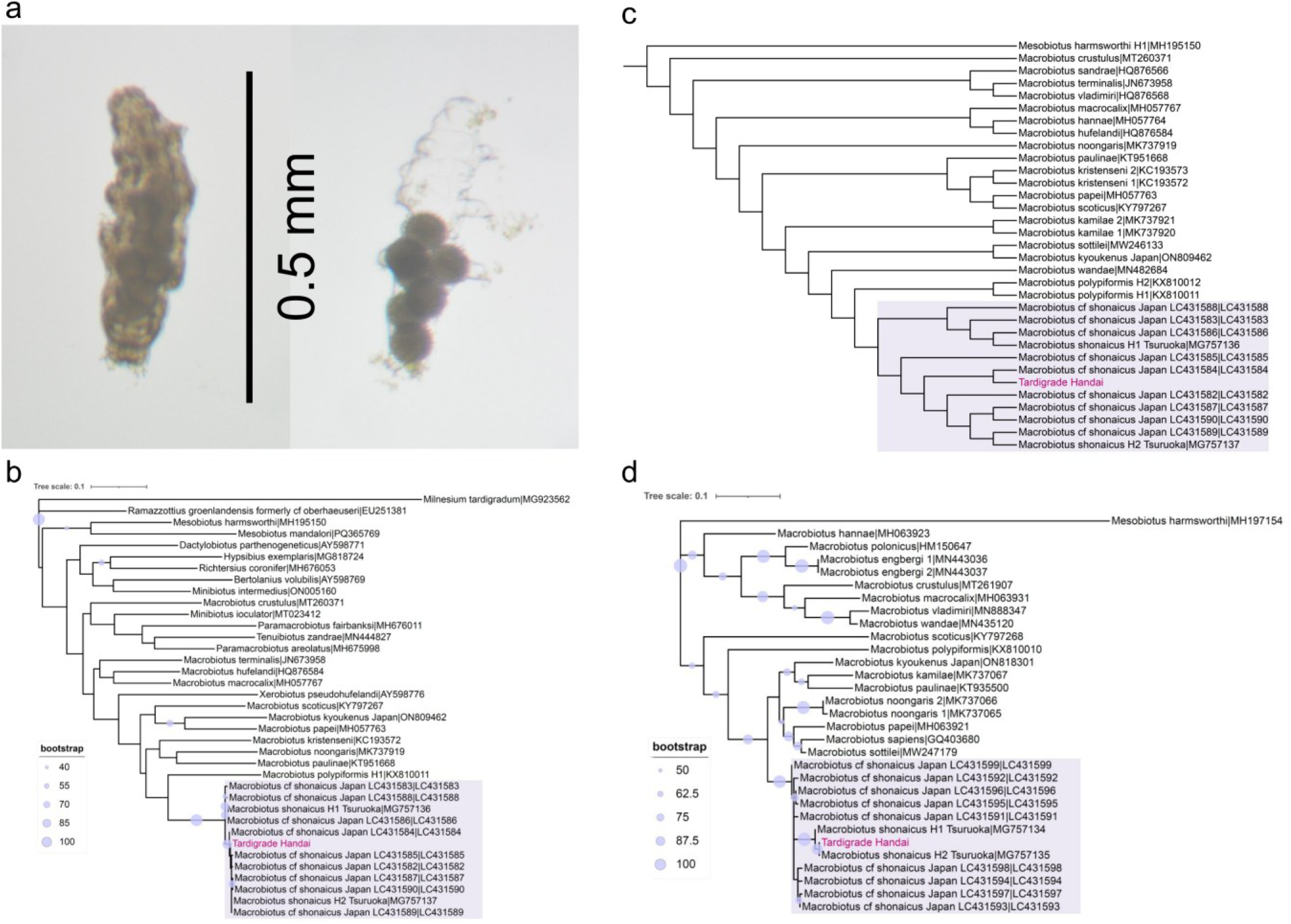
Cultured tardigrades and molecular species assignment. a, Light micrographs of an animal containing eggs before oviposition (left) and eggs within a shed cuticle (right) from the laboratory culture. Scale bar, 0.5 mm. b, Broad COI maximum-likelihood phylogeny including representatives of Macrobiotidae and selected other eutardigrade groups, rooted with *Milnesium tardigradum*. c, Focused COI phylogeny of *Macrobiotus shonaicus* and related species, rooted with *Mesobiotus harmsworthi*. d, Focused ITS2 maximum-likelihood phylogeny, rooted with *Mesobiotus harmsworthi*. The culture analyzed in this study is labeled “Tardigrade Handai” in magenta, and the clade containing *M. shonaicus* is shaded. GenBank accession numbers are included in the tip labels. Trees were inferred under the GTR+I+G4 model with 1,000 bootstrap replicates. Circle sizes indicate the displayed bootstrap support values in b and d; tree scale bars indicate substitutions per site.

**Supplementary Fig. 2.**
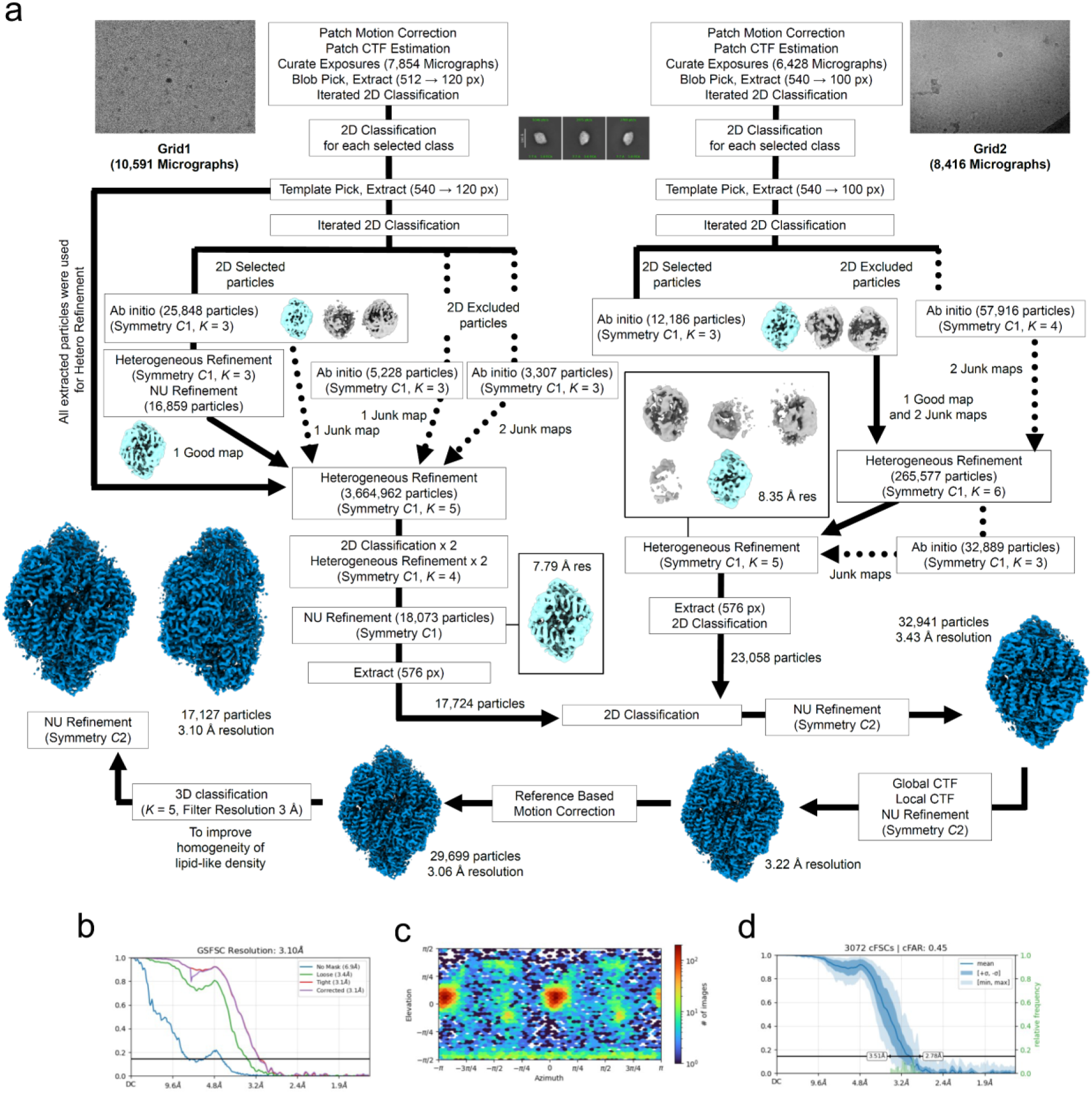
Cryo-EM image processing and reconstruction of *Ms*Vg6. a, Processing workflow for particles from two grids prepared from the same crude extract. The datasets were processed separately before selected particles were combined. Ab initio reconstruction and heterogeneous refinement were performed without imposed symmetry (*C*1), followed by non-uniform refinement with *C*2 symmetry, global and local CTF refinement, and reference-based motion correction. An intermediate reconstruction of the combined dataset reached 3.06 Å resolution from 29,699 particles. Further three-dimensional classification to improve the homogeneity of lipid-like density yielded a subset of 17,127 particles and the final 3.10-Å reconstruction used for all *Ms*Vg6 map visualizations in the figures. Particle numbers, symmetries and intermediate resolutions are indicated. b, Gold-standard Fourier shell correlation (FSC) curves for the 3.10-Å reconstruction. c, Particle viewing-direction distribution. d, Directional conical FSC analysis. NU, non-uniform.

**Supplementary Fig. 3.**
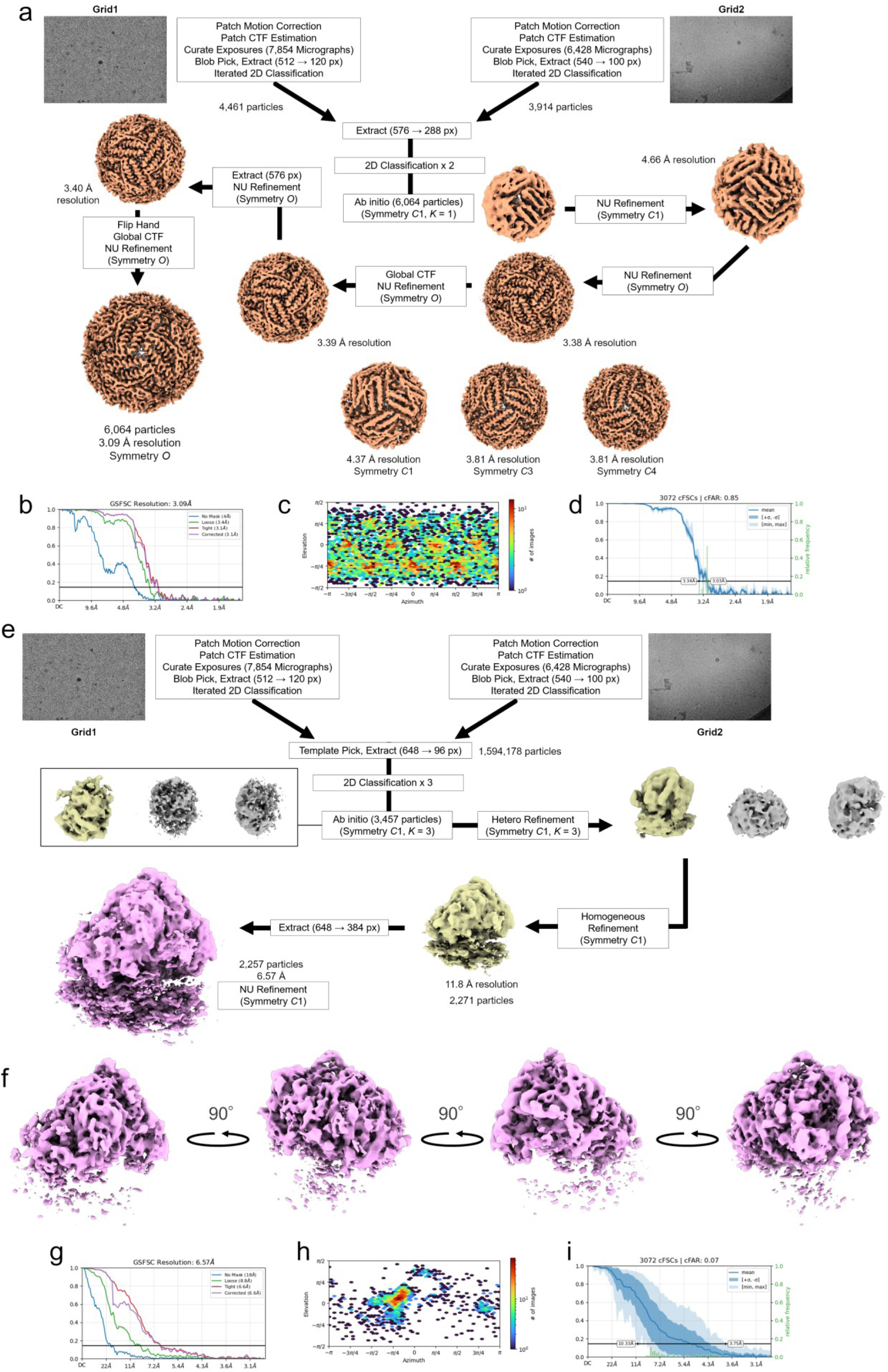
Cryo-EM image processing and reconstruction of *Ms*Ftn and the 80S ribosome. a, *Ms*Ftn processing workflow, showing combination of particles from two grids, two-dimensional classification, ab initio reconstruction, non-uniform refinement, CTF refinement and correction of map handedness. The final reconstruction with octahedral (*O*) symmetry reached 3.09 Å resolution from 6,064 particles. Reconstructions with *C*1, *C*3 and *C*4 symmetry are also shown. b–d, Gold-standard FSC curves, particle viewing-direction distribution and directional conical FSC analysis for *Ms*Ftn, respectively. e, Ribosome processing workflow, including template-based picking, two-dimensional classification, ab initio reconstruction, heterogeneous and homogeneous refinement, and final non-uniform refinement with *C*1 symmetry. The final reconstruction reached 6.57 Å resolution from 2,257 particles. f, Ribosome map displayed at a high contour level to emphasize the 60S subunit, in four views related by successive 90° rotations. g–i, Gold-standard FSC curves, particle viewing-direction distribution and directional conical FSC analysis for the ribosome, respectively. NU, non-uniform.

**Supplementary Fig. 4.**
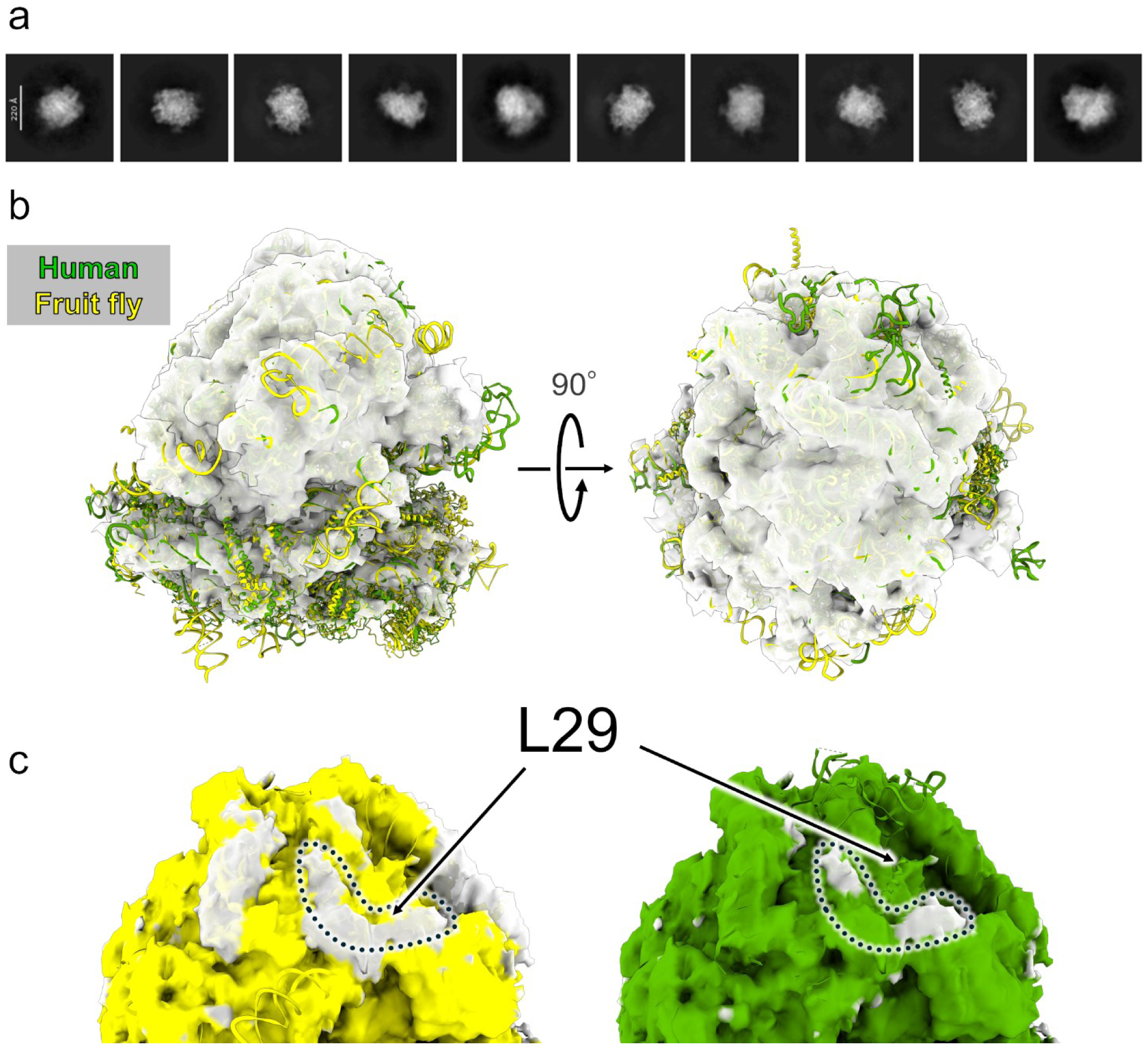
Cryo-EM reconstruction of the endogenous tardigrade 80S ribosome. a, Representative two-dimensional class averages of ribosome particles. Scale bar, 220 Å. b, The 6.57-Å cryo-EM map shown in two views related by a 90° rotation, with the 60S portions of fitted 80S reference models from human (green, PDB 4UG0)^14^ and *Drosophila* (yellow, PDB 4V6W)^15^. c, Close-up comparisons with the *Drosophila* (left) and human (right) reference structures. Map regions within 8 Å of each fitted reference model are colored according to the model. Dotted outlines highlight density adjacent to ribosomal protein L29 that lies outside the 8-Å coloring regions for both fitted reference models. This density was left unassigned.

**Supplementary Fig. 5.**
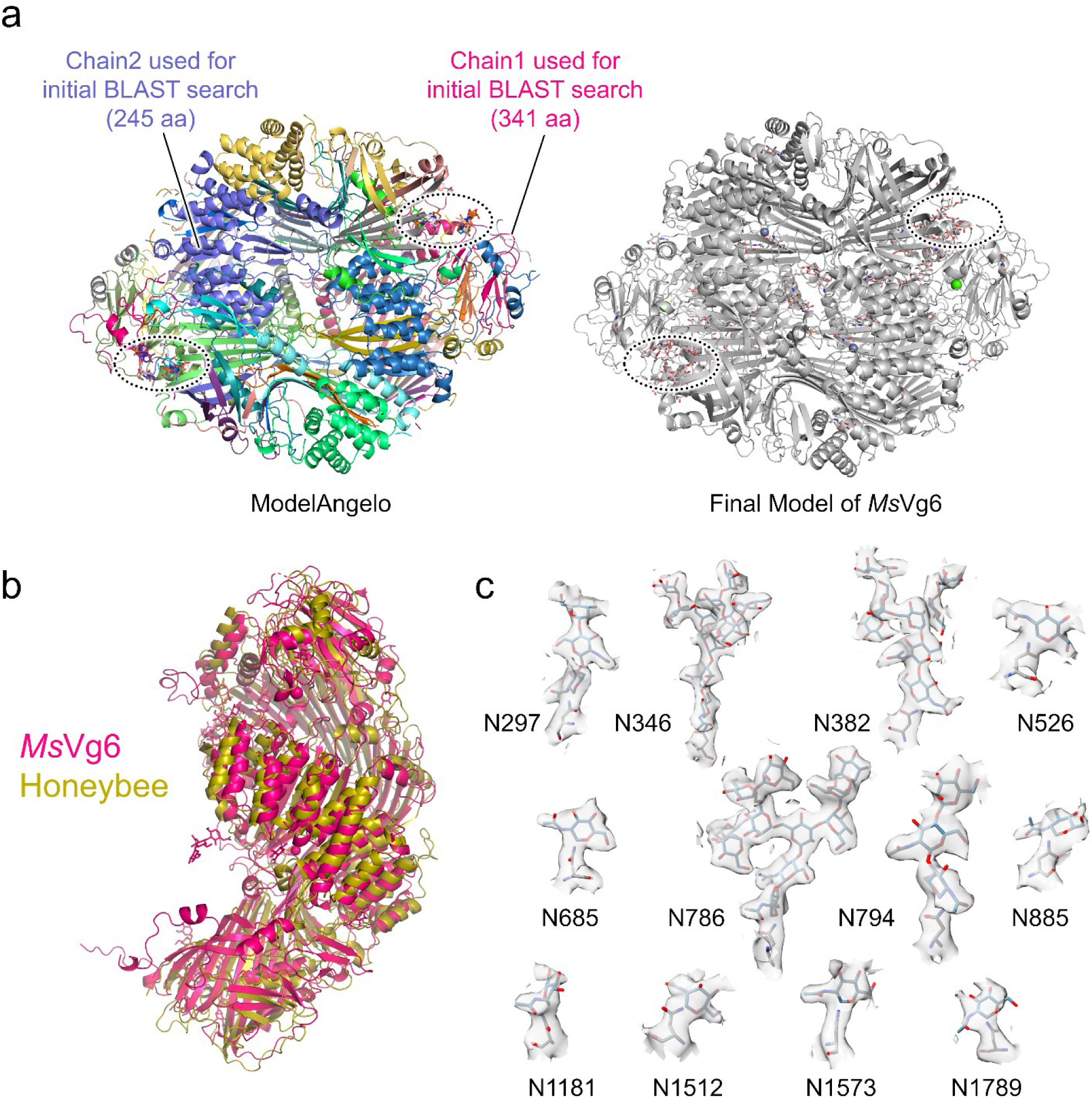
*Ms*Vg6 model assignment, comparison with honeybee Vg and N-glycan densities. a, The initial ModelAngelo model is colored by chain (left), and the final model is shown in gray with glycans as sticks (right). The chains used for the initial BLAST search are indicated. Dotted outlines mark regions where ModelAngelo placed nucleic acid fragments and N-linked glycans were modeled in the final structure. b, Superposition of an *Ms*Vg6 protomer (magenta) and honeybee Vg (PDB 9ENR; olive). c, Local cryo-EM densities for N-linked glycans at Asn297, Asn346, Asn382, Asn526, Asn685, Asn786, Asn794, Asn885, Asn1181, Asn1512, Asn1573 and Asn1789. Glycans and the attached asparagine residues are shown as sticks within semitransparent gray density surfaces. Carbon, oxygen and nitrogen atoms are colored blue-gray, red and blue, respectively.

**Supplementary Fig. 6.**
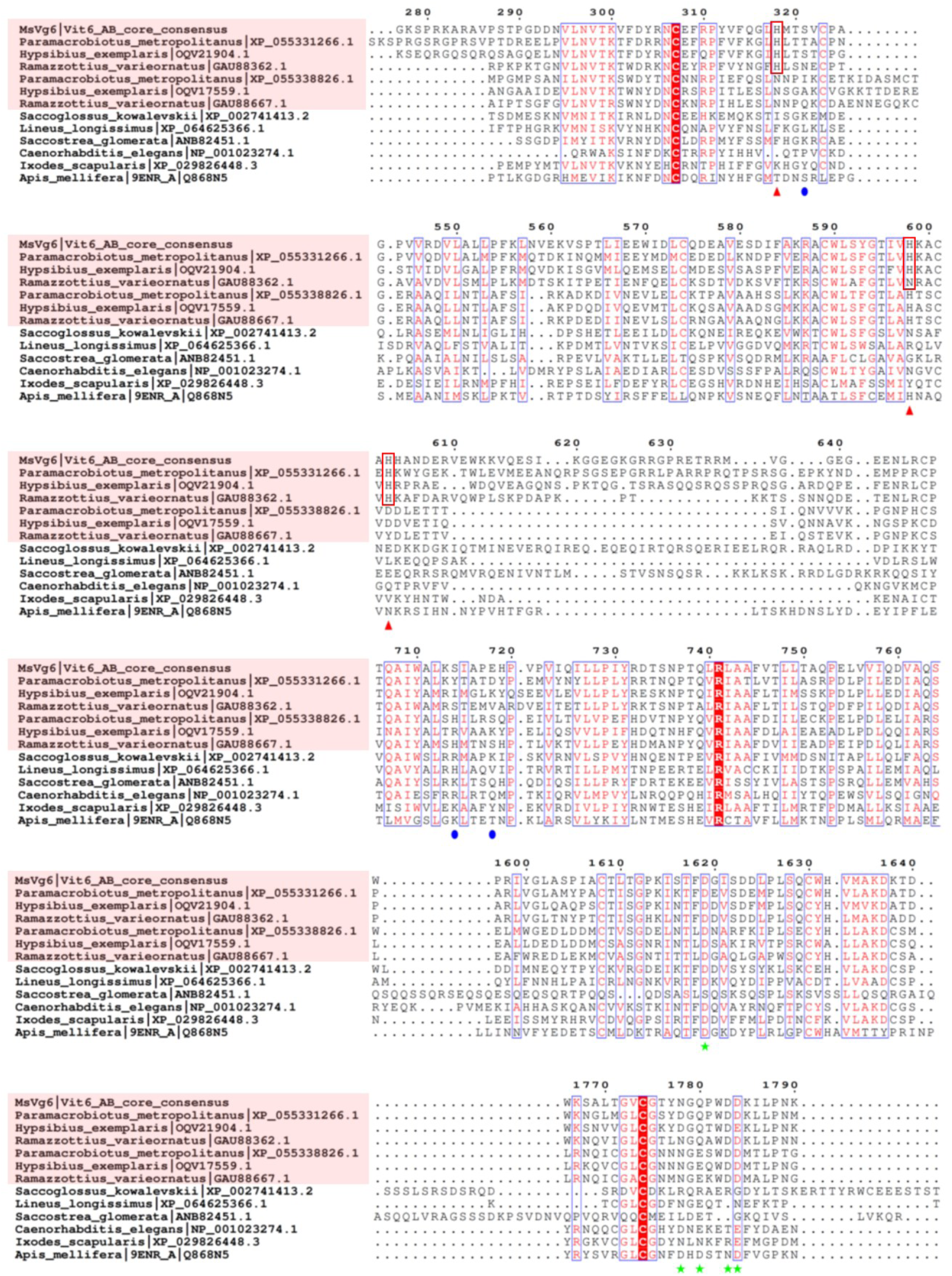
Sequence alignment of *Ms*Vg6 and representative Vg homologues. Selected regions of an alignment of *Ms*Vg6 and 12 homologues are shown, including six tardigrade homologues and honey bee Vg (PDB 9ENR; UniProt Q868N5). Tardigrade sequence labels are shaded pink. Numbering follows *Ms*Vg6. Red triangles mark His318, His598 and His603 at the putative Zn-binding site; red outlines highlight the corresponding positions in *Ms*Vg6 and its three closest homologues. Blue circles mark nearby Ser321, Ser714 and Glu718. Green stars mark Asp1619, Asn1778, Gln1780, Asp1783 and Asp1784 at the putative Ca-binding site. White letters on red backgrounds indicate identical residues, and red letters and blue boxes indicate conserved similarities. Dots denote alignment gaps.

**Supplementary Fig. 7.**
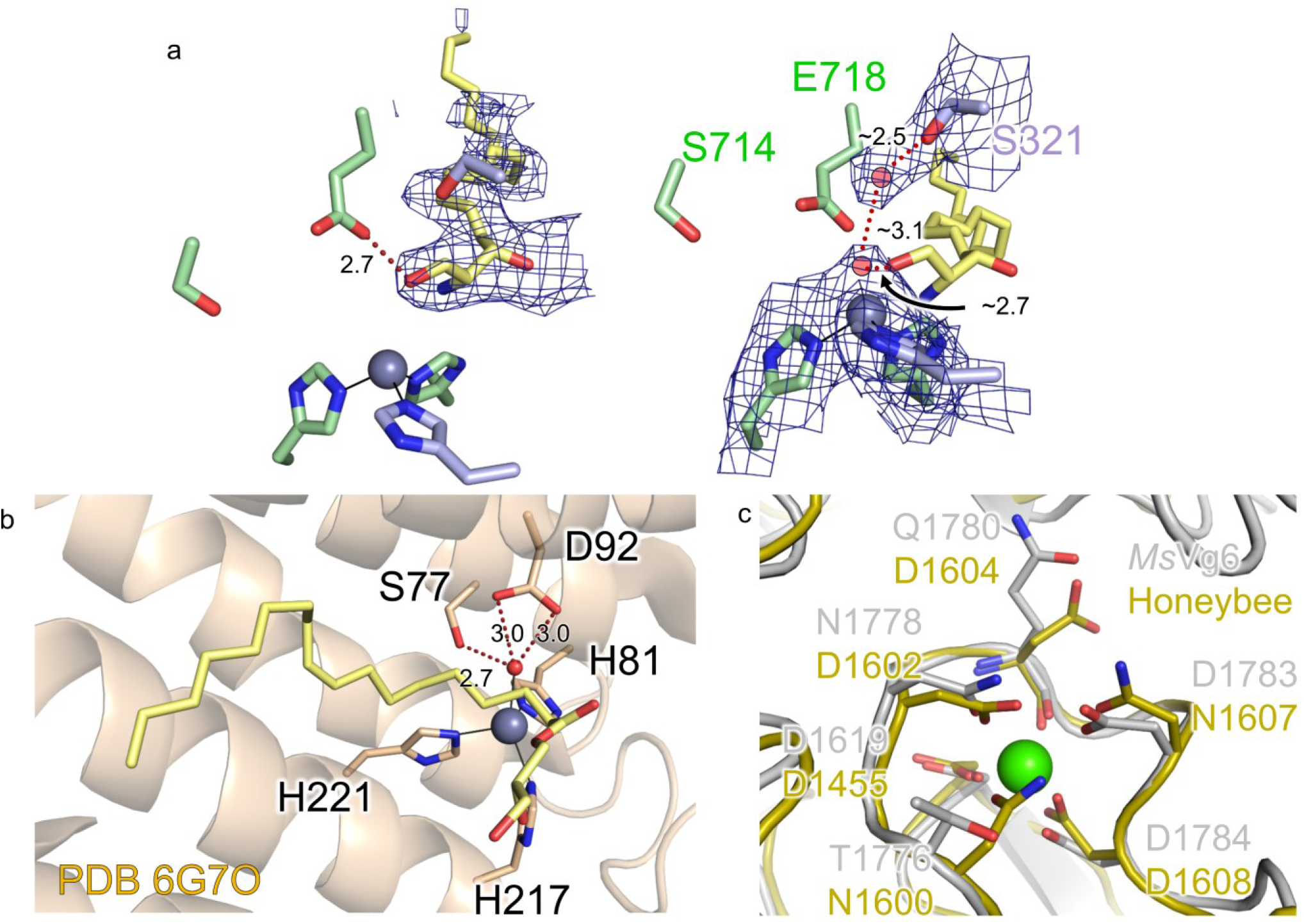
Comparisons of the putative metal-binding environments of *Ms*Vg6 with ACER3 and honeybee Vg. a, Two views of the putative Zn site and adjacent elongated density in *Ms*Vg6, with a tentative sphingosine-like fit shown in yellow. Protein residues are shown as sticks, the putative Zn ion as a sphere, and cryo-EM density as blue mesh. Glu718, Ser321 and Ser714 are labeled in the right-hand view; red spheres indicate putative solvent molecules. b, The Zn-binding site of human ACER3 (PDB 6G7O),^27^ showing the coordinating residues His81, His217 and His221, the Zn-bound water molecule, and nearby Ser77 and Asp92. The monoolein molecule (OLB) modeled in the reference structure is shown in yellow. Dotted lines and distances in Å indicate selected local contacts. The comparison highlights similarities in the local metal-binding environments; however, it does not establish the chemical identity of the *Ms*Vg6 ligand or ceramidase activity. c, Superposition of the local vWD structures of *Ms*Vg6 (gray) and honeybee Vg (PDB 9ENR; olive). Selected side chains are shown as sticks, with oxygen and nitrogen atoms colored red and blue, respectively. The green sphere represents the putative Ca²⁺ ion modeled in *Ms*Vg6. The comparison shows a similar local arrangement of oxygen-containing side chains at the site corresponding to the Ca²⁺-binding site predicted in honeybee Vg. Sequence conservation at the corresponding positions is shown in Supplementary Fig. 6.

**Supplementary Fig. 8.**
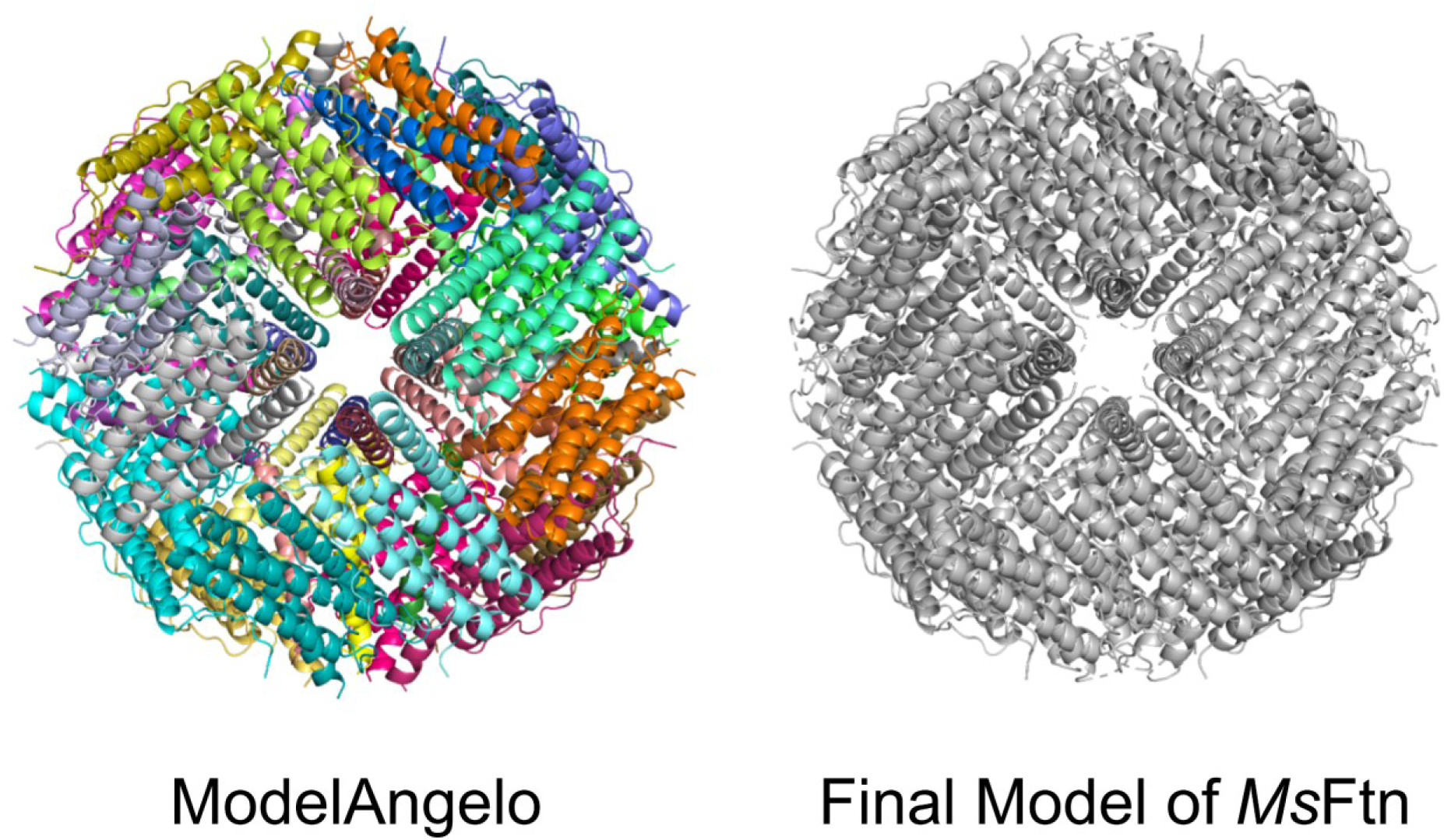
Comparison of the initial ModelAngelo model and the final *Ms*Ftn model. The initial model is colored by chain (left), and the final model is shown in gray (right).

**Supplementary Fig. 9.**
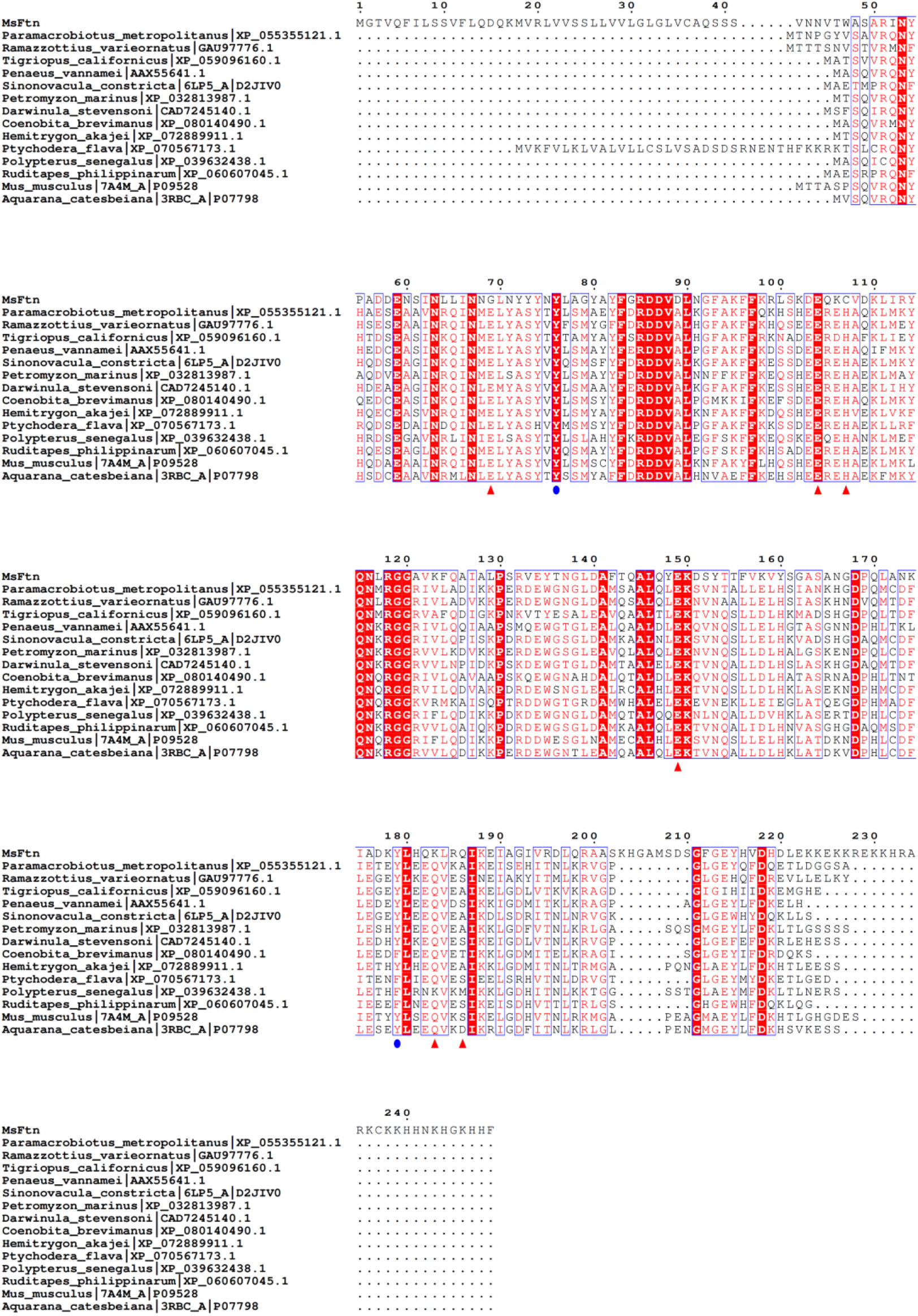
Sequence alignment of *Ms*Ftn and representative ferritin homologues. The alignment includes *Ms*Ftn and 14 homologues, including mouse heavy-chain ferritin (PDB 7A4M; UniProt P09528) and bullfrog M ferritin (PDB 3RBC; UniProt P07798). Species names and sequence accession numbers are indicated. Numbering above the alignment follows *Ms*Ftn. White letters on red backgrounds indicate identical residues, and red letters and blue boxes indicate conserved similarities. Dots denote alignment gaps. Red triangles mark Gly69, Glu104, Cys107, Glu149, Lys183 and Gln186 of *Ms*Ftn, corresponding to metal-coordinating positions in the bullfrog ferritin reference. Blue circles mark Tyr76 and Tyr179, corresponding to Tyr30 and Tyr133 in bullfrog ferritin. The alignment highlights substitutions at positions corresponding to the canonical ferroxidase center and the extended C-terminal sequence of *Ms*Ftn.

**Supplementary Fig. 10.**
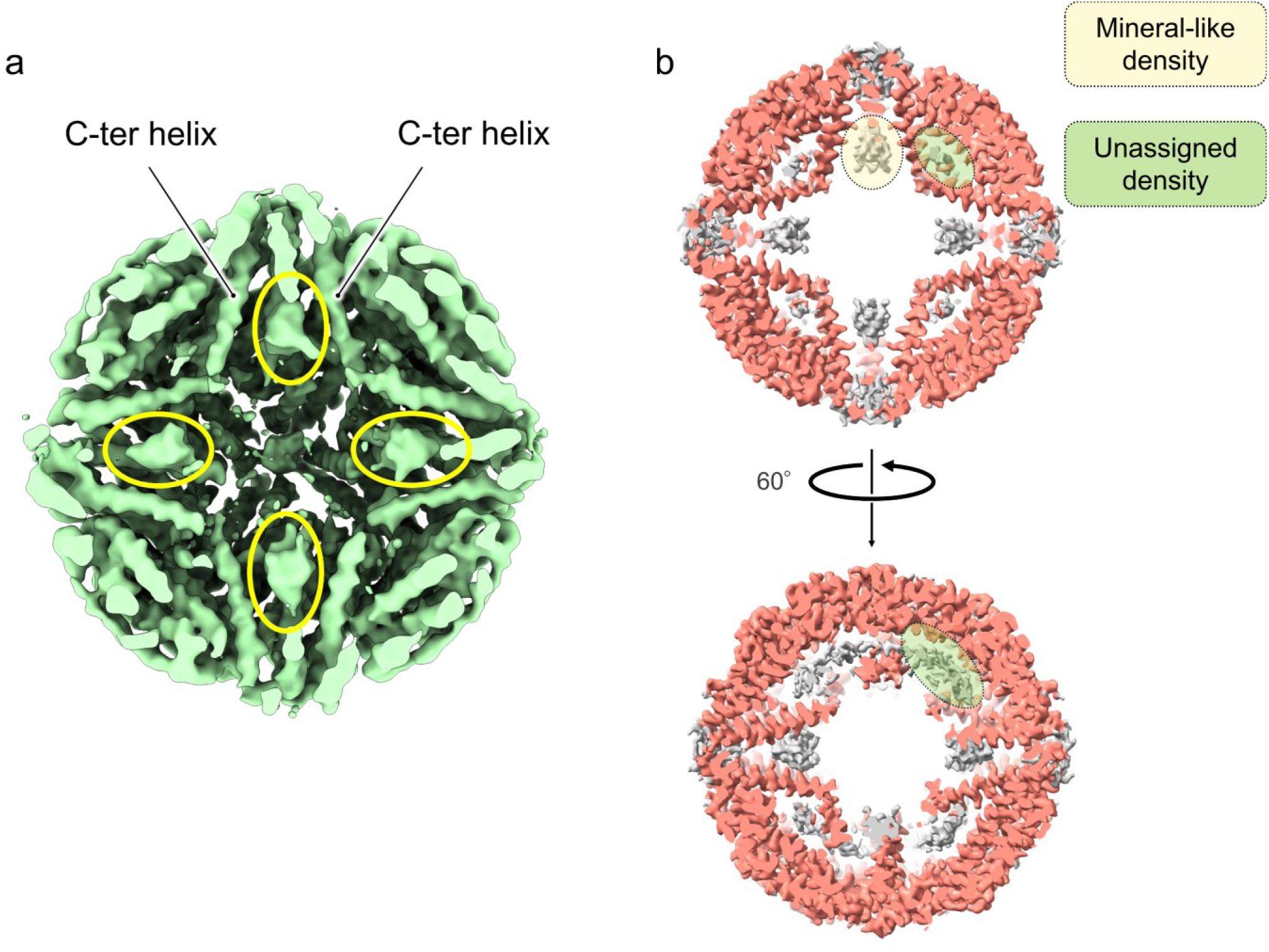
Internal non-protein densities in the *Ms*Ftn cage. a, Sectioned map reconstructed without imposed symmetry (*C*1). Yellow outlines highlight internal mineral-like densities, and the positions of C-terminal helices are indicated. b, Sectioned maps reconstructed with octahedral (O) symmetry, shown in two views related by a 60° rotation. Dotted outlines and pale shading distinguish the mineral-like density beneath the fourfold-axis region (yellow) from the additional unassigned density along the inner wall (green). Neither density was assigned a definitive chemical composition.

**Supplementary Table 1.**
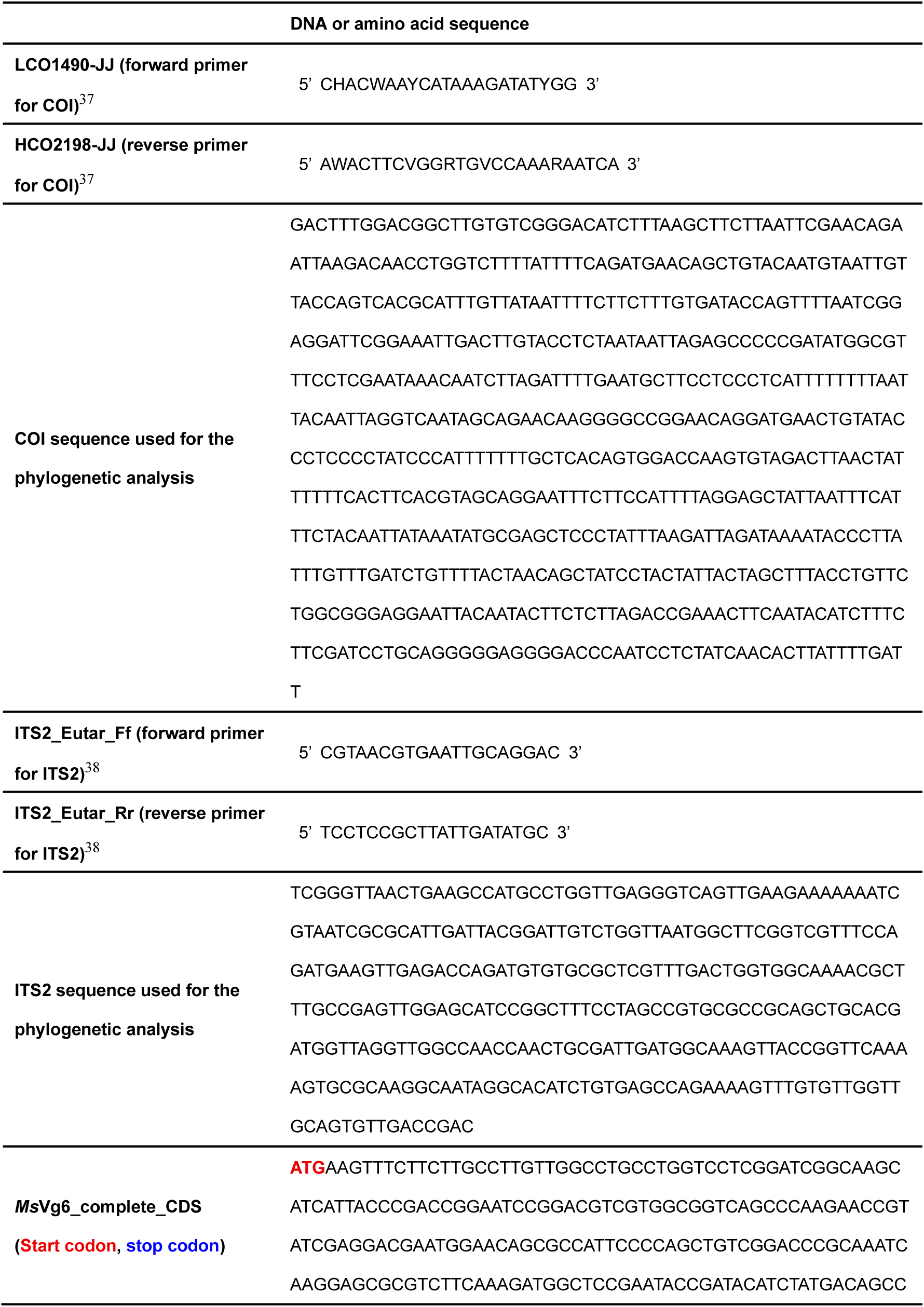
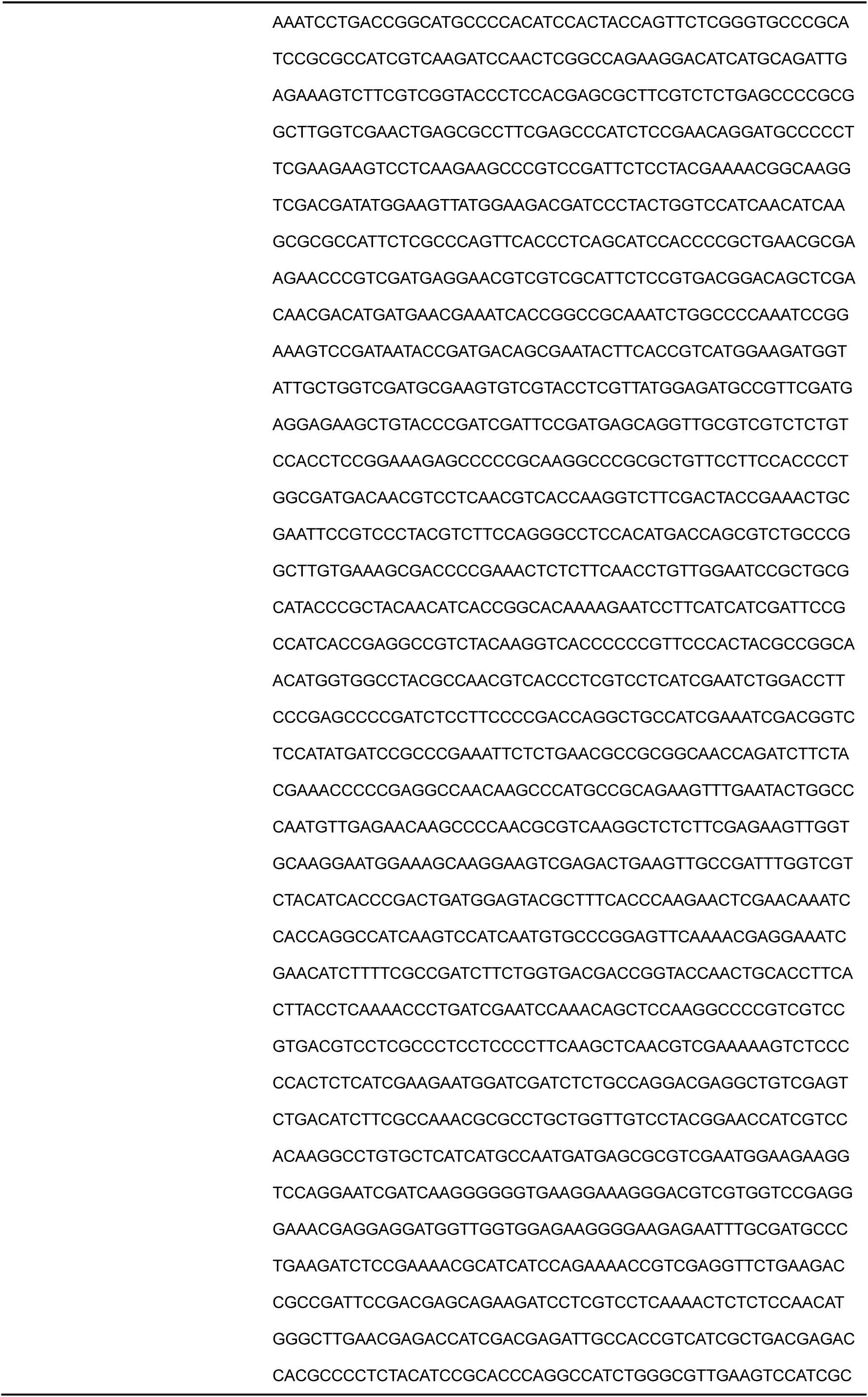
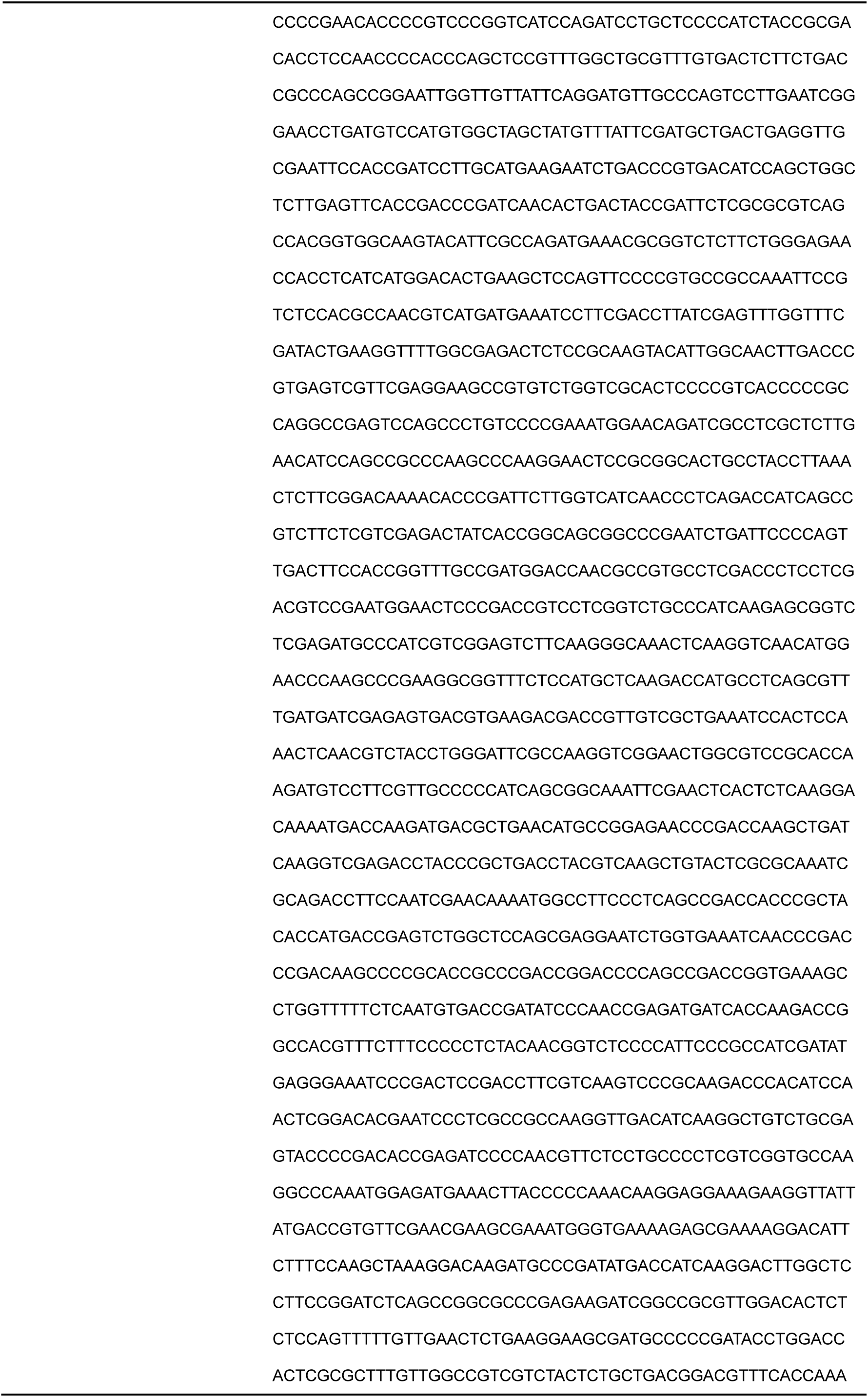
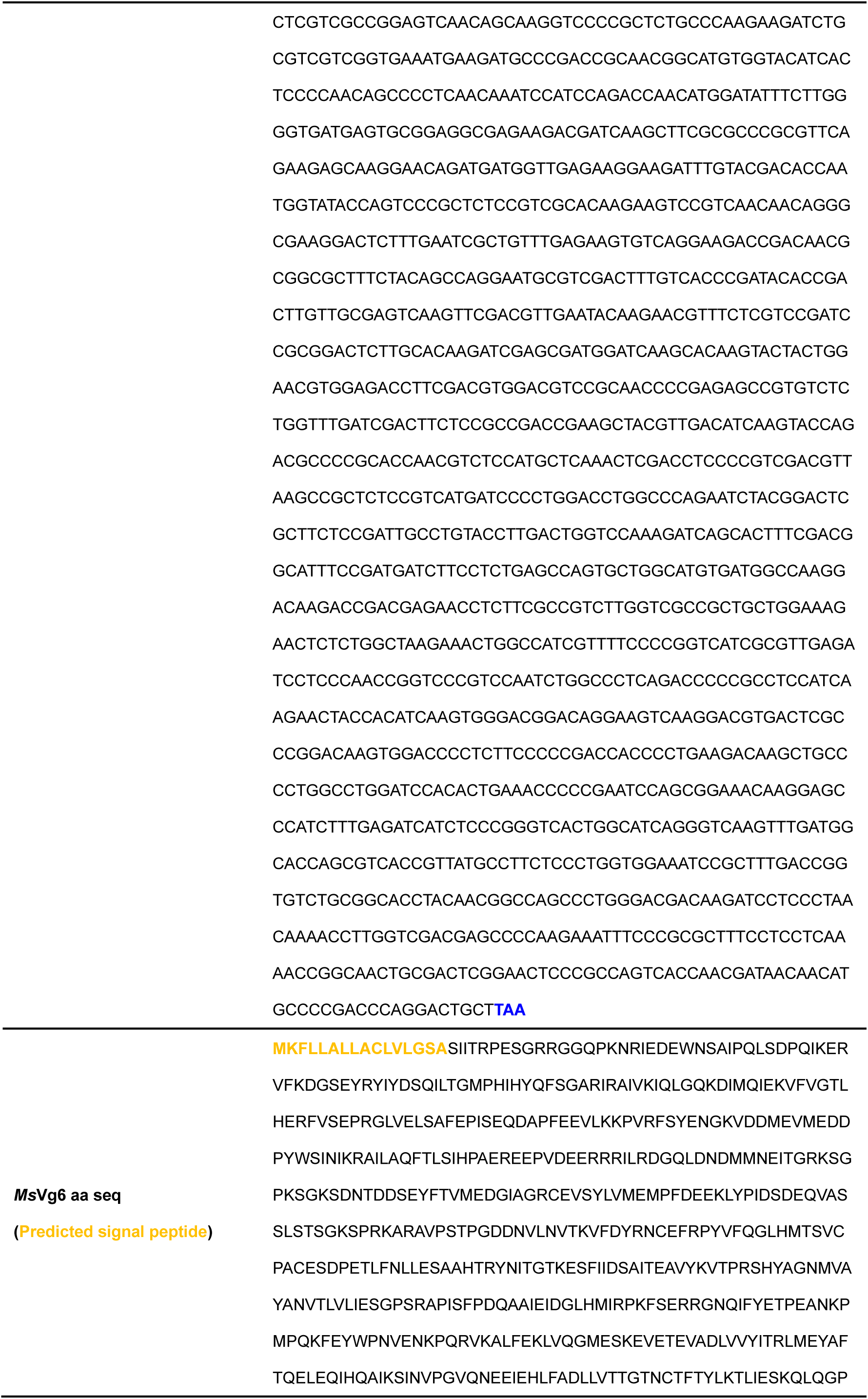
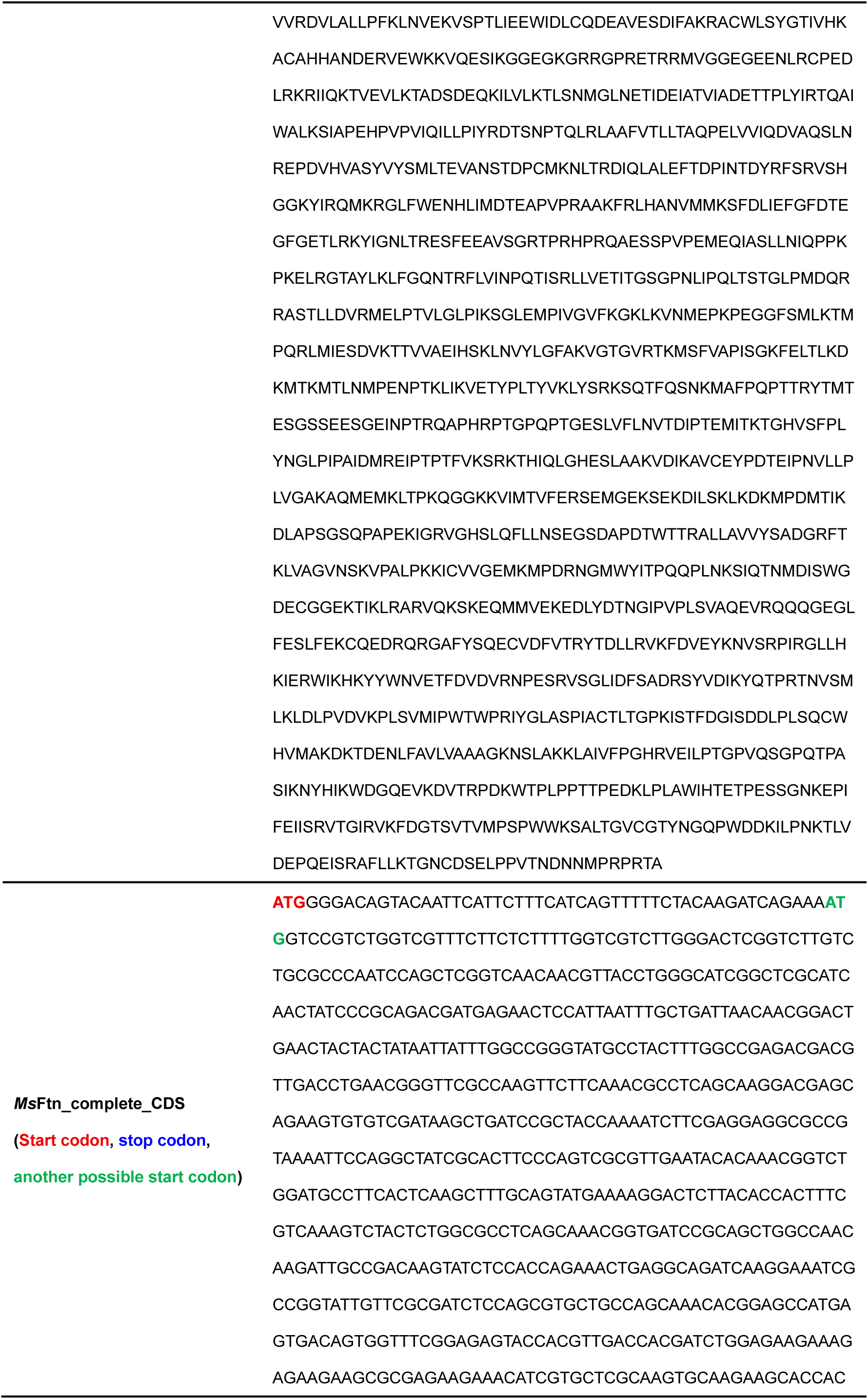
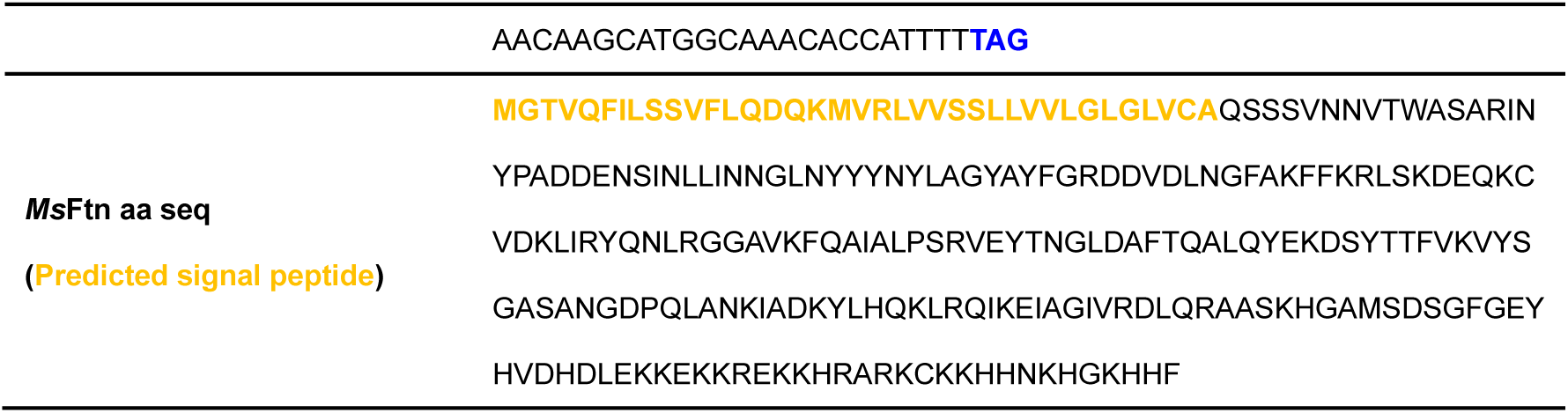
Primers used for COI and ITS2 amplification and coding and amino acid sequences of *Ms*Vg6 and *Ms*Ftn. Predicted signal peptides identified using SignalP 6.0^51^ are highlighted in orange in the amino acid sequences. In the *Ms*Ftn coding sequence, the upstream ATG used for residue numbering is shown in red, the downstream candidate initiation ATG (corresponding to Met18) in green and the stop codon in blue. *Ms*Ftn residue numbering starts at the upstream candidate initiation codon; the actual translation-initiation site remains undetermined.

**Supplementary Table 2.** Cryo-EM data collection, reconstruction and model refinement statistics.

|  | ***MsVg6*** | ***MsFtn*** | **Ribosome** |
| --- | --- | --- | --- |
| Extended PDB ID | pdb_000046QY | pdb_000046QZ | - |
| EMDB ID | EMD-83988 | EMD-83989 | EMD-83990 |
| **Data collection and processing** |  |  |  |
| Magnification | 60,000 | 60,000 | 60,000 |
| Voltage (kV) | 200 | 200 | 200 |
| Electron exposure (e-/Å<sup>2</sup>) | 40.0 | 40.0 | 40.0 |
| Defocus range (μm) | -0.7 to -1.7 | -0.7 to -1.7 | -0.7 to -1.7 |
| Pixel size (Å) | 0.83 | 0.83 | 0.83 |
| Symmetry imposed | C2 | O | C1 |
| Initial particle images (no.) | 3,930,535 | 2,050,898 | 1,594,178 |
| Final particle images (no.) | 17,127 | 6,064 | 2,257 |
| Map resolution (Å) | 3.10 | 3.09 | 6.57 |
| FSC threshold | 0.143 | 0.143 | 0.143 |
| **Refinement** |  |  |  |
| Initial model used | ModelAngelo | ModelAngelo |  |
| Model resolution (Å) | 3.4 | 3.3 |  |
| FSC threshold | 0.5 | 0.5 |  |
| Map sharpening *B*-factor (Å<sup>2</sup>) | 61.2 | 76.2 |  |
| Model composition |  |  |  |
| Non-hydrogen atoms | 14,084 | 1,518 |  |
| Protein residues | 13,339 | 1,518 |  |
| Ligands | 745 | - |  |
| *B*-factors (Å<sup>2</sup>) |  |  |  |
| Protein | 49.6 | 52.5 |  |
| Ligand | 52.0 | - |  |
| R.m.s. deviations |  |  |  |
| Bond lengths (Å) | 0.006 | 0.004 |  |
| Bond angles (°) | 0.938 | 0.851 |  |
| Validation |  |  |  |
| MolProbity score | 1.76 | 1.38 |  |
| Clashscore | 8.73 | 6.95 |  |
| Ramachandran plot (%) |  |  |  |
| Favored | 95.85 | 98.34 |  |
| Allowed | 3.97 | 1.66 |  |
| Outliers | 0.18 | 0.00 |  |
| Model vs Map |  |  |  |
| CC (mask) | 0.85 | 0.85 |  |
| CC (main chain) | 0.84 | 0.84 |  |
| CC (side chain) | 0.81 | 0.77 |  |

**Supplementary Table 3.**
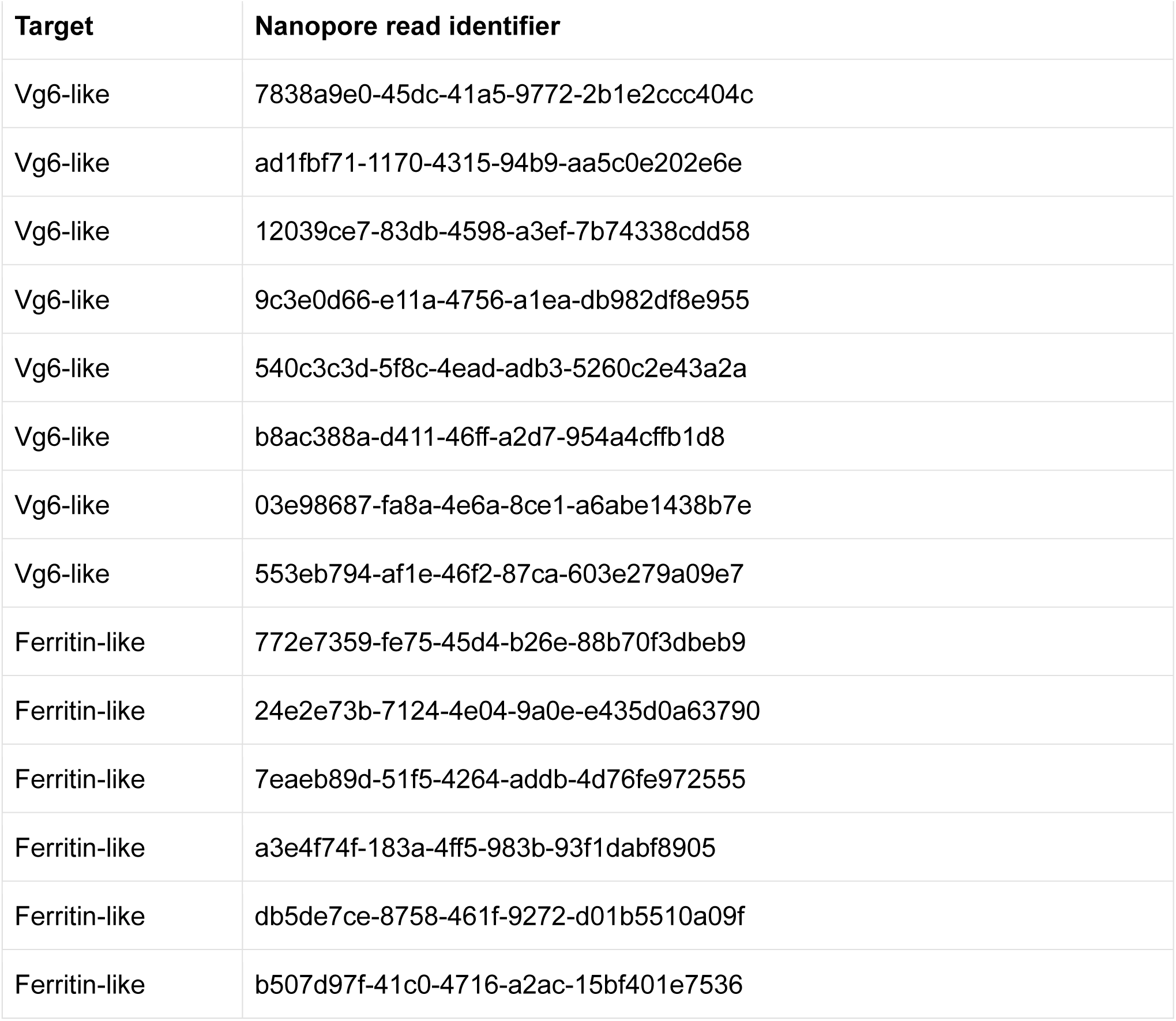
Long-read cDNA reads used for consensus construction. Each identifier is the unique read ID in the Nanopore cDNA sequencing dataset and identifies an individual sequencing read used to construct the Vg6-like or ferritin-like consensus. These are read-level identifiers, not gene or database accession numbers. The table lists eight Vg6-like reads and six ferritin-like reads.

